# Cell Surface SARS-CoV-2 Nucleocapsid Protein Modulates Innate and Adaptive Immunity

**DOI:** 10.1101/2021.12.10.472169

**Authors:** Alberto Domingo López-Muñoz, Ivan Kosik, Jaroslav Holly, Jonathan W. Yewdell

**Affiliations:** Cellular Biology Section, Laboratory of Viral Diseases, NIAID (NIH), Bethesda, Maryland, United States

## Abstract

SARS-CoV-2 nucleocapsid protein (N) induces strong antibody and T cell responses. Although considered to be localized in the cytosol, we readily detect N on the surface of live cells. N released by SARS-CoV-2 infected cells or N-expressing transfected cells binds to neighboring cells by electrostatic high-affinity binding to heparan sulfate and heparin, but not other sulfated glycosaminoglycans. N binds with high affinity to 11 human chemokines, including CXCL12β, whose chemotaxis of leukocytes is inhibited by N from SARS-CoV-2, SARS-CoV-1, and MERS CoV. Anti-N Abs bound to the surface of N expressing cells activate Fc receptor-expressing cells. Our findings indicate that cell surface N manipulates innate immunity by sequestering chemokines and can be targeted by Fc expressing innate immune cells. This, in combination with its conserved antigenicity among human CoVs, advances its candidacy for vaccines that induce cross-reactive B and T cell immunity to SARS-CoV-2 variants and other human CoVs, including novel zoonotic strains.

## INTRODUCTION

Despite the unprecedented expeditious development and deployment of highly effective vaccines, the rapid selection of SARS Coronavirus (CoV) 2 (SARS-CoV-2) spike glycoprotein (S) antibody (Ab) escape mutants threatens to delay the return to pre-pandemic conditions. To broaden vaccination and reduce SARS-CoV-2 related acute and chronic disease, it is crucial to improve our knowledge of innate and adaptive immunity to CoVs.

CoVs encode four major structural proteins. S, membrane (M), and envelope (E) proteins are localized in the viral surface envelope. N binds to viral RNA through electrostatic interactions, forming cytoplasmic helical nucleocapsids that associate with M to enable virus budding into early secretory compartments. As the most abundantly expressed SARS-CoV-2 protein, N induces strong Ab and T_CD8+_ immune responses^1,2^. Although CoV N is widely considered to be strictly localized in the cytoplasm, cell surface expression of RNA viruses N is more the rule than the exception. Early studies with monoclonal Abs (mAbs) reported surface expression of influenza A and vesicular stomatitis virus N ^3,4^. Influenza N is a target for Ab-complement-mediated cell lysis^3^, Ab redirected T cell lysis^5^, and is targeted by protective Abs in mice^6^. N and N-like RNA genome binding proteins are expressed on the surface of cells infected with other human viruses, including measles^7^, respiratory syncytial^8^, lymphocytic choriomeningitis^9^, and human immunodeficiency virus^10^.

Here, we examine the expression of human CoV N on the cell surface and its participation in innate and adaptive immunity.

## RESULTS

### SARS-CoV-2 N is robustly expressed on the infected cell surface

We examined cell surface expression of SARS-CoV-2 N by imaging Vero cells 24 h post-infection (hpi) with wild-type (wt) or a recombinant SARS-CoV-2 expressing eGFP (SARS-CoV-2_eGFP). To exclusively detect cell surface N, we incubated live cells with primary and fluorophore-conjugated secondary antibodies at 4°C prior to fixation and mounting for confocal imaging. This releveled clear surface N staining over mock-infected (mock) background levels, using S or eGFP as markers of infected cells (Fig. 1a, maximum intensity projection images of z-stack). We similarly found N on the surface of BHK-21_humanACE2(hACE2), Caco-2, Calu-3, CHO-K1_hACE2, and HEK293-FT_hACE2 cells infected with wt or eGFP SARS-CoV-2 at 24 hpi (Extended Data Fig. 1, 2). Depending on the cell type, we observed a variable degree of colocalization between N and S, particularly remarkable in Vero (Fig. 1a), Calu-3, CHO-K1_hACE2, and HEK293-FT_hACE2 cells (Extended Data Fig. 1). We noted a dramatic syncytia formation in hACE2 overexpressing BHK-21_hACE2 and HEK293-FT_hACE2 cells as reported ^11^.

**Fig. 1:**
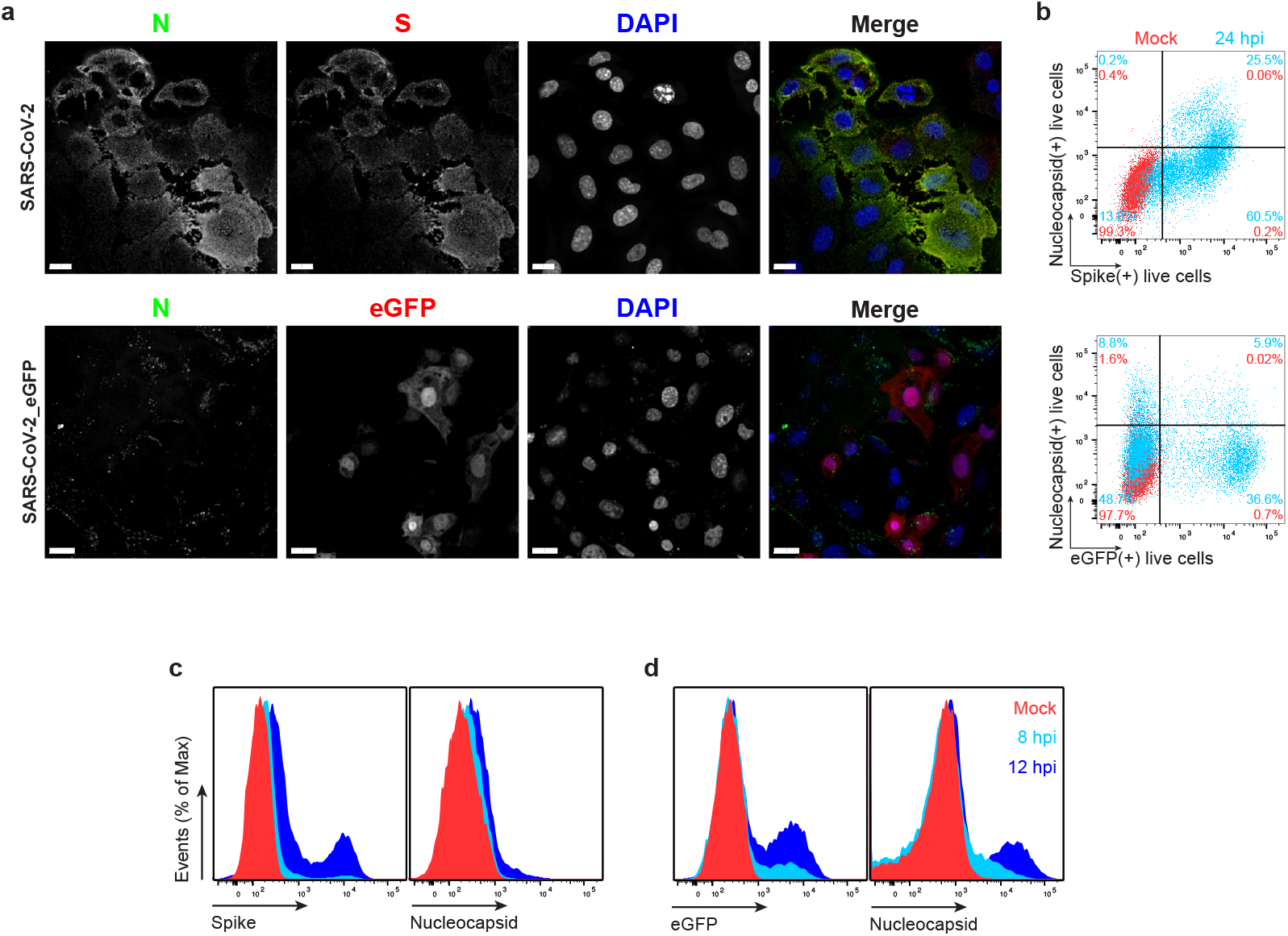
SARS-CoV-2 N is expressed on the surface of live cells early during infection. **a,** Maximum intensity projections (MIP) of laser confocal microscopy z-stack images of infected Vero cells with wild-type SARS-CoV-2 (top panels) or SARS-CoV-2_eGFP, stained live at 24 hpi (MOI = 1). Scale bar = 20 μm. Images are representative of at least three independent experiments with similar results. **b,** Flow cytometry analyses of Vero cells inoculated with wild-type (top) or eGFP expressing (bottom) SARS-CoV-2 (MOI = 1), stained live at 24 hpi against SARS-CoV-2 S and N proteins. Representative dot plots of flow cytometry analyses showing double staining of surface S and N, and eGFP proteins, indicating the percentage of the gated cell population for each quadrant of the double staining. Data are representative of at least three independent experiments, each performed with triplicate samples. **c, d,** Time course of surface S, N, and eGFP proteins expression in live infected Vero cells with wild-type (**c**) and eGFP reporter (**d**) SARS-CoV-2 at 8 and 12 hpi (MOI = 1). Representative histogram overlays of surface S, N, and intracellular eGFP proteins of flow cytometry analyses. Data are representative of one experiment out of at least two independent experiments performed in triplicate.

To measure N surface expression more quantitatively, we performed flow cytometry analyses of live infected cells 24 hpi. Surface N was detected on a subpopulation of S or eGFP expressing cells for each of the seven cell types examined (Fig. 1b, Extended Data Fig. 1–3). N was also detected on the surface of live cells infected with Alpha (B.1.1.7), Beta (B.1.351) and Delta (B.1.617.2) SARS-CoV-2 variants (Extended Data Fig. 4). Via flow cytometry, we determined the kinetics of N expression on the surface of Vero (Fig. 1c-d), BHK-21_hACE2, and A549_hACE2 cells (Extended Data Fig. 5). As early as 8 hpi, we observed a significant surface signal for N protein in live Vero and BHK-21_hACE2 infected cells, while it took slightly longer (12 hpi) for A549_hACE2 cells (Extended Data Fig. 5). Notably, depending on cell type and marker of infection (S vs. eGFP) we detected cells expressing N but not S or eGFP on a fraction of cells, ranging from less than 1% to 43% (Extended Data Fig. 1, 2). This is consistent with several mechanisms acting alone or in combination: differential expression of SARS-CoV-2 gene products in individual cells ^12^, complete retention of S in the secretory pathway ^13^, and transfer of N from infected to non-infected cells.

To determine whether N cell surface expression requires other SARS-CoV-2 gene products, we transfected cells with an expression plasmid encoding N. Staining of live BHK-21, CHO-K1 or HEK293-FT cells transfectants revealed up to 7-fold more N mAb binding over background levels obtained with cells transfected with a control plasmid expressing eGFP (Fig. 2a). N-surface expression increased between 24 and 72 h post-transfection, providing further evidence for the specificity of staining, and demonstrating that cell surface expression is an intrinsic property of biosynthesized N.

**Fig. 2:**
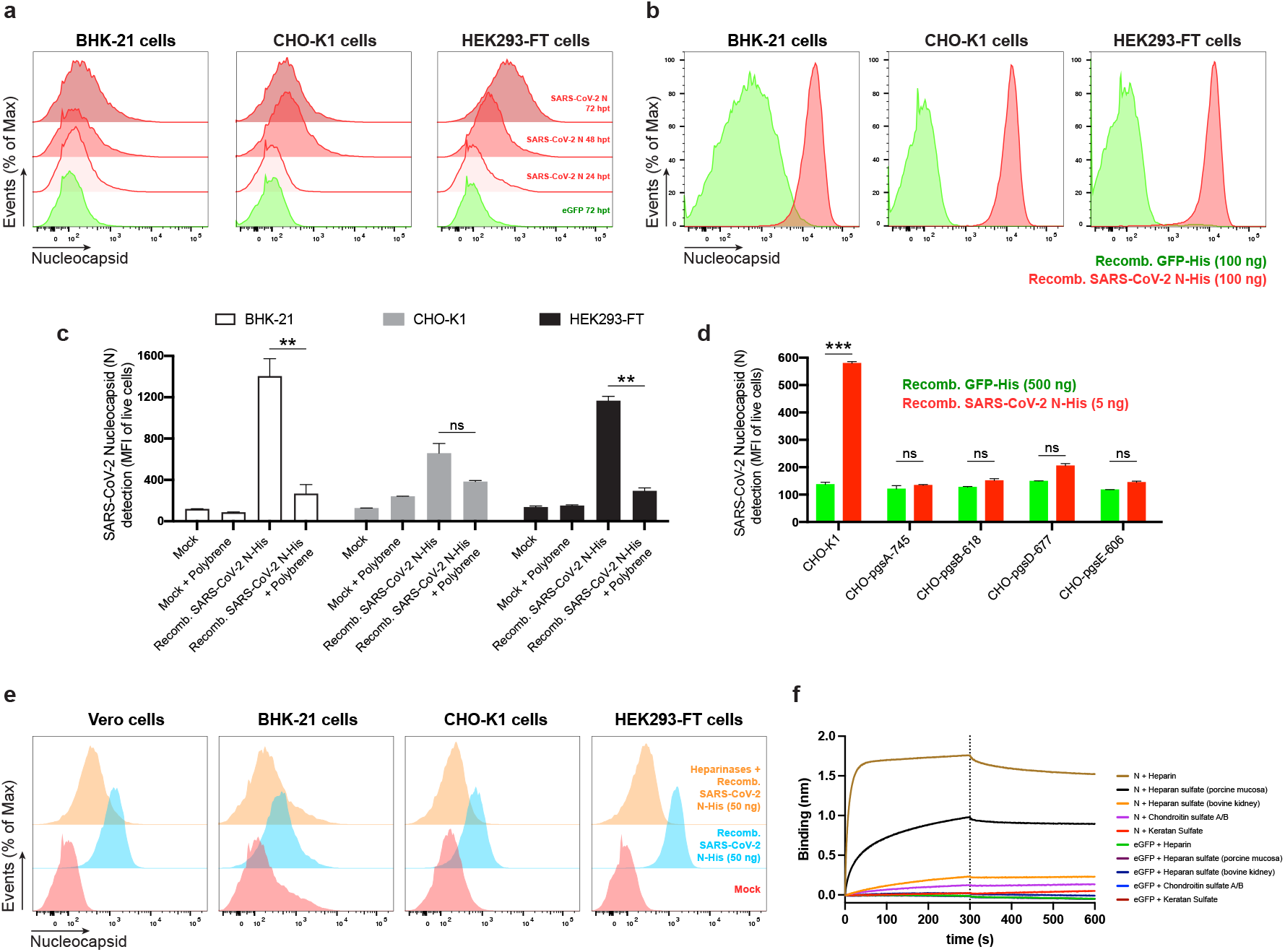
SARS-CoV-2 N cell surface binding is independent of other viral genes and is specifically mediated by heparan sulfate/heparin. **a,** Histogram semi-overlays of kinetics (24-72 h) of surface N protein expression in BHK-21, CHO-K1 and HEK293-FT cells transiently transfected with a plasmid encoding eGFP or N protein, detected with Abs by flow cytometry. **b,** Histogram overlays of analyses of exogenous rN binding to BHK-21, CHO-K1 and HEK293-FT cells, incubated with recombinant eGFP or N protein for 15 min, washed twice, stained live with Abs, and analyzed by flow cytometry. **c,** Electric charge neutralization assays with a cationic polymer (polybrene). BHK-21, CHO-K1 and HEK293-FT cells were incubated with 50 ng of rN protein for 15 min, washed twice, incubated with 10 μg/ml of polybrene, washed twice again, stained live with Abs, and analyzed by flow. **d,** Different GAG-deficient CHO cells were incubated with recombinant eGFP or N protein for 15 min, washed twice, stained live with Abs, and analyzed by flow cytometry. **e,** Heparinase treatment completely abrogates the cell ability to bind and retain N protein. Flow cytometry histogram semi-overlays of BHK-21, CHO-K1 and HEK293-FT cells treated with heparinases for 1 h, washed twice, incubated with 50 ng of rN protein for 15 min, washed twice again, stained live with Abs, and analyzed. **f,** BLI sensorgrams from binding assays of sulfated GAGs to immobilized N or eGFP proteins. Streptavidin-coated biosensors were loaded with equivalent amounts of N or eGFP, measuring their ability to bind each GAG. Sensorgrams show association and dissociation phases, where the vertical dotted line indicates the end of the association step. In **c, d,** the mean fluorescent intensity (MFI) of detected surface N protein from live cells is plotted, showing mean +/-SEM (n = 2). Student’s two-tailed unpaired *t*-test was used to compare N-incubated cells vs. N-incubated and polybrene treated cells (**c**), and to compare GFP-vs. N-incubated cells (**d**): *ns* (statistically nonsignificant) *p* > 0.01, ** *p* < 0.01, *** *p* < 0.001. The analyses were repeated with different protein preparations, and one representative assay out of at least three independent assays performed in duplicate is shown.

### Exogenous N binds to cells

To examine whether N surface expression requires its synthesis in the cell, we incubated BHK-21, CHO-K1, or HEK293-FT cells with exogenous purified recombinant N (rN) for 15 min at 37 C°resulted in. This resulted in strong flow cytometry staining (2-log shift) with anti-N mAbs relative to control cells incubated with an irrelevant protein (Fig. 2b).

N interacts with negatively charged viral RNA through highly positively charged RNA-binding domains^14,15^. We examined charge-based N binding to the cell surface by treating rN coated cells with polybrene, a cationic polymer that neutralizes surface electrostatic charges. Flow cytometric analysis showed that polybrene decreases rN bound to live BHK-21, CHO-K1, and HEK293-FT cells to similar levels, with the magnitude of the effect proportional to the amount of bound N (Fig. 2c).

For most mammalian cells, glycosaminoglycans (GAGs) are the major negatively charged molecule on the plasma membrane^16^. To assess the contribution of GAGs to N cell surface binding, we used a panel of GAG-deficient CHO cells^17^. Each of the GAG-deficient cell lines tested failed to bind rN over levels observed with recombinant GFP (Fig. 2d). The panel included cells with complete absence of GAGs (CHO-pgsA-745, HO-pgsB-618), and cells with defects in synthesizing heparan sulfate and heparin but no other GAGs (CHO-pgsD-677, CHO-pgsE-606). Consistent with these findings, treating Vero, BHK-21, CHO-K1, or HEK293-FT cells with heparinase I, II, and III in combination, to depolymerize heparan sulfate/heparin polysaccharide chains to disaccharides, cells dramatically reduced binding of exogenous rN (Fig. 2e, Extended Data Fig. 6c). We directly confirmed N binding to heparin by using biolayer interferometry (BLI), where we directly demonstrate N specific nanomolar affinity binding to heparan sulfate and heparin, but not to other sulfated GAGs (Fig. 2f, Extended Data Fig. 6d, e).

Together, these findings indicate that N binds to the cell surface by interacting with heparan sulfate and heparin in a charge-dependent manner.

### N is transferred from expressing to non-expressing cells

In SARS-CoV-2 24 hpi immunofluorescence and flow cytometry experiments (Fig. 1a, b, Extended Data Fig. 1, 2), we observed cells expressing N but not S or GFP as early as 8 and 12 hpi (Extended Data Fig. 5), with increasing numbers of cells over time after infection (Extended Data Fig. 7). To determine whether N can be transferred from infected to non-infected cells, we added SARS-CoV-2 to a co-culture of infectable (hACE2 expressing) and non-infectable (non-hACE2 expressing) CHO-K1 cells (we confirmed that hACE2 is required for infecting CHO cells with SARS-CoV-2, see Supplementary Fig. 1a). We pre-stained non-infectable cells with CellTrace™ Violet to enable unambiguous flow identification after co-culture (Supplementary Fig. 1b). Remarkably, co-cultured non-ACE2 expressing uninfected CHO-K1 cells (also confirmed by the absence of S expression) have a higher cell surface N signal than infected cells (Fig. 3a, Supplementary Fig. 1c). N transfer from infected cells required GAG expression on uninfected cells, as shown by near background staining by GAG-deficient CHO cells (Fig. 3b, c, Supplementary Fig. 1d, e). N is also transferred from HEK293-FT or BHK-21 cells transiently expressing N from a transgene to co-cultured un-transfected cells (Fig. 3d).

**Fig. 3:**
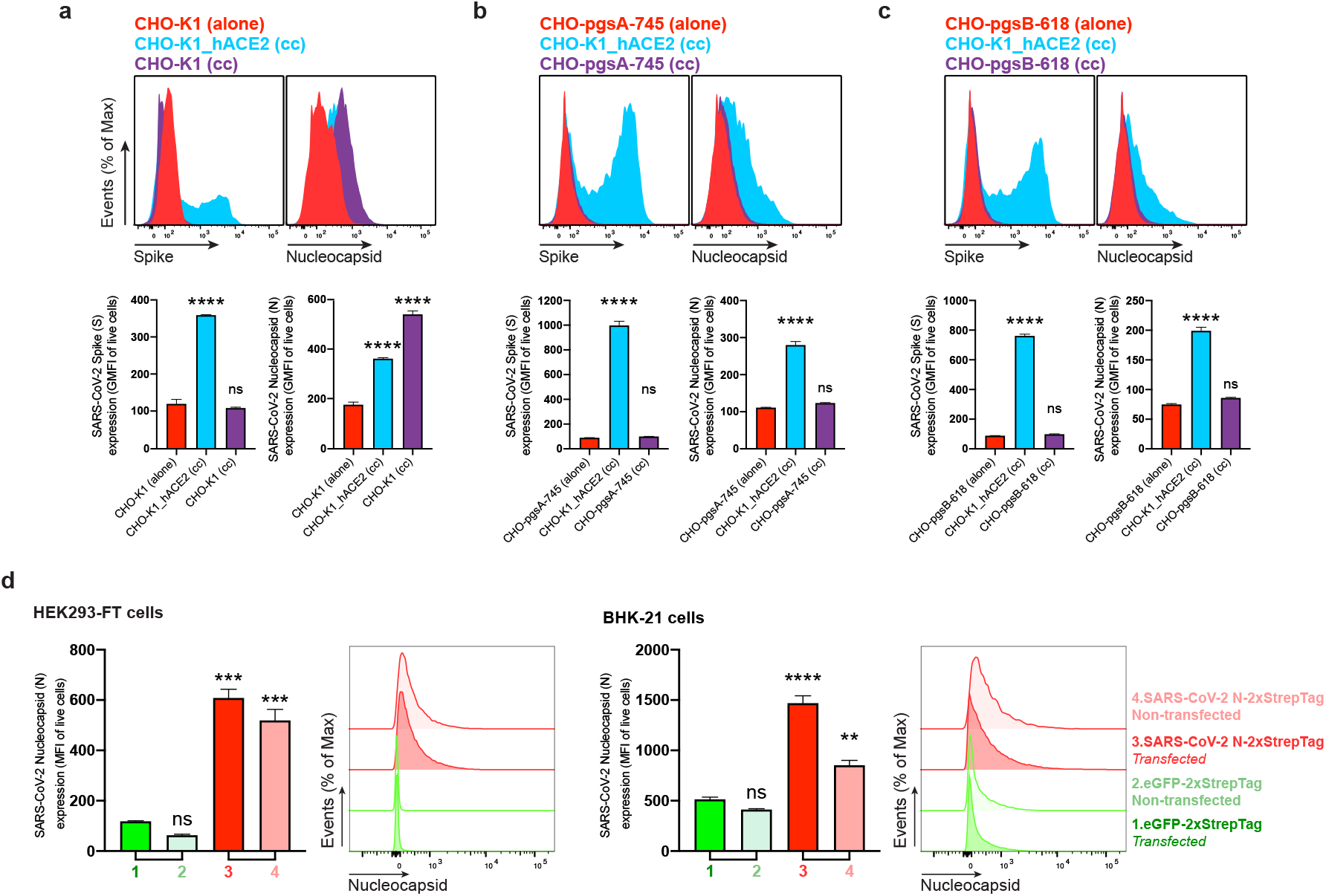
SARS-CoV-2 N protein is transferred to the cell surface of neighboring uninfected cells. Flow cytometry analyses of N transfer assays between donor and recipient co-cultured cells. **a-c,** N transfer assays between infectable and non-infectable co-cultured cells. CHO-K1 (**a**), GAG-deficient CHO-pgsA-745 (**b**) and CHO-pgsB-618 (**c**) cells (non-infectable), alone or co-cultured (cc) with CHO-K1_hACE2 cells (infectable), were inoculated with SARS-CoV-2 (MOI = 1) and stained live with Abs at 24 hpi against surface SARS-CoV-2 S and N proteins. Non-infectable cells were stained with CellTraceTM Violet prior co-culture with infectable cells (**Supplementary Fig. 1**). For dot plots showing double staining of surface S and N with percentages of the gated cell population for each quadrant see **Supplementary Fig. 1c-e**. **d,** N transfer assays between transfected and non-transfected co-cultured cells. HEK293-FT and BHK-21 cells were transiently transfected with a plasmid encoding eGFP or N protein. After 24 h, non-transfected HEK293-FT or BHK-21 cells were stained with CellTraceTM Violet prior to be added and co-cultured with their transfected counterparts. Cells were stained live after 24 h with Abs and analyzed. For each assay, the following is shown: histogram overlays and semi-overlays of surface N and S proteins, as well as the GMFI or MFI geometric MFI (GMFI) of live cells expressing S and N proteins is plotted, showing mean +/- SEM (n = 3). One representative experiment of at least three independent experiments performed in triplicate is shown. One-way ANOVA and Dunnett’s Multiple comparison test were used to compare all conditions against non-infectable cells cultured alone within each assay, or against eGFP-transfected cells: *ns* (nonsignificant) *p* > 0.05, ** *p* < 0.01, *** *p* < 0.001, **** *p* < 0.0001.

Based on these findings, we conclude that N protein synthesis leads to its release from cells and its robust transfer to non-synthesizing cells, where it is retained by binding heparin/heparan sulfate.

### SARS-CoV-2 N inhibits chemokine function but enables Ab based immune cell activation

The robust expression of N on the surface of infected and surrounding cells suggests a significant evolutionary function. SARS-CoV-2, like most viruses, induces the release of pro-inflammatory cytokines by infected cells. Could N interfere with this signaling? We examined the ability of immobilized N to interact with 64 human cytokines by BLI. Remarkably N bound CCL5, CCL11, CCL21, CCL26, CCL28, CXCL4, CXCL9, CXCL10, CXCL11, CXCL12β, and CXCL14 chemokines with micromolar to nanomolar affinities (Fig. 4a, Extended Data Fig. 8). By contrast, none of the other SARS-CoV-2 immobilized structural (S, M, E) or non-structural (ORFs 3a, 3b, 6, 7a, 7b, 8, 9b, 9c, and 10) proteins tested interacted with any of the cytokines in the panel with affinities higher than that observed for immobilized GFP (Extended Data Fig. 9a). Kinetic curves of N binding to each chemokine were biphasic, deviating from first-order binding (1:1) and consistent with heterogeneous binding (Extended Data Fig. 8).

**Fig. 4:**
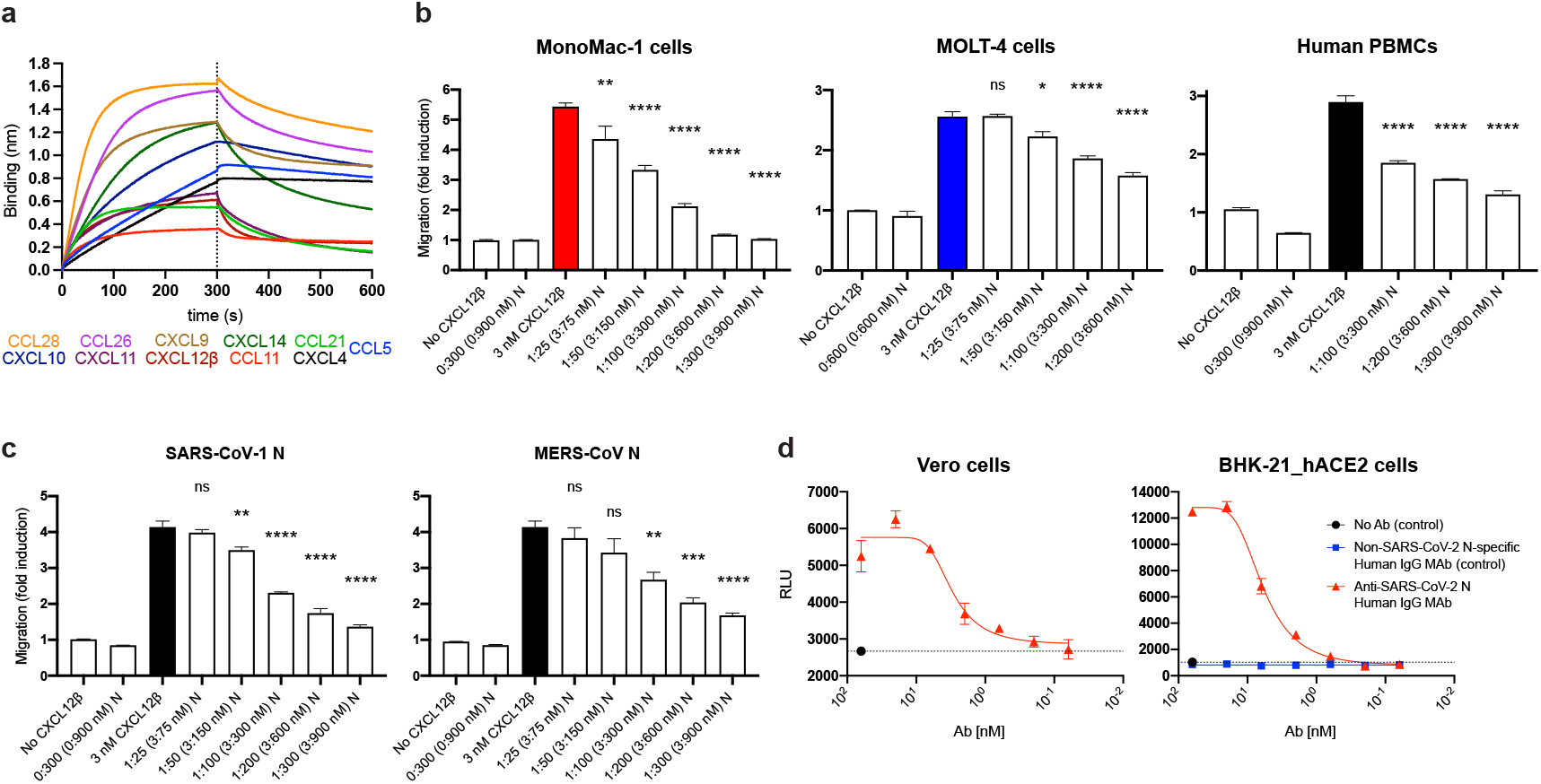
SARS-CoV-2 N protein modulates innate and adaptive immunity, **a, b,** N binds chemokines through the GAG-binding domain of chemokines and inhibits in vitro chemokine-mediated leukocyte migration. **a,** BLI sensorgrams of binding assays showing association and dissociation phases of the interaction between N protein and 11 positively bound chemokines at a concentration of 100 nM out of 64 human cytokines tested. The dotted line indicates the end of the association step. The analyses were repeated with different purified rN protein preparations. One representative assay of three independent assays is shown. **b,** SARS-CoV-2 N blocks CXCL12β chemotaxis of MonoMac-1 cells, MOLT-4 cells and human PBMCs. **c,** SARS-CoV-1 N and MERS-CoV N block CXCL12β chemotaxis of MonoMac-1 cells. CXCL12β was incubated alone or in the presence of the indicated viral protein in the lower chamber of transwell migration devices. Migrated cells from the top chamber were detected in the lower chamber at the end of the experiment. The induction of migration shows means +/- SEM (n = 3) from one representative assay performed in triplicate out of at least three independent assays. One-way ANOVA and Dunnett’s Multiple comparison test were used to compare all conditions (except no chemokine and viral protein alone conditions) against migration induced by chemokine alone (colored bar): *ns* (nonsignificant) *p* > 0.05, * *p* < 0.05, ** *p* < 0.01, *** *p* < 0.001, **** *p* < 0.0001. **d,** N protein is a target for Ab-based immunity. ADCC reporter bioassays were performed on SARS-CoV-2-infected Vero and BHK-21_hACE2 target cells (24 hpi, MOI =1) using decreasing concentrations of a human mAb against the N protein and Jurkat effector reporter cells. After overnight incubation, luciferase expression to gauge cell activation was measured. Data show mean +/- SEM (n = 3) of one representative assay out of three independent experiments performed in triplicate. Dashed lines show background signal detected in the absence of Ab.

Chemokine function is based on interaction with both surface GAGs and their specific receptors located in the surface of leukocytes. We found that blocking the GAG-binding domain of chemokines by increasing concentrations of heparan sulfate (from bovine kidney) and chondroitin sulfate A and B, N binding to its subset of chemokines was abrogated (Extended Data Fig. 10). This indicates that N binds to chemokines through their GAG-binding domain.

What is the functional relevance of N-chemokine interactions? Transwell chemotaxis experiments with monocyte-like cells (MonoMac-1), T lymphocyte-like cells (MOLT-4), and human healthy donor PBMCs revealed that rN blocked CXCL12β-induced chemotaxis in a concentrationdependent manner (Fig. 4b). While rN from different vendors showed comparable results inhibiting CXCL12β-mediated induction of migration of MonoMac-1 cells, the S1 protein (subunit 1 of the S protein, containing the receptor-binding domain) had no inhibitory effect on CXCL12β-induced chemotaxis (Supplementary Fig. 2). Extending these findings, rN from both SARS-CoV-1 and MERS-CoV also inhibited CXCL12β induced migration of MonoMac-1 cells (Fig. 4c).

mAbs to mouse CoV N have been reported to exert anti-viral activity *in vitro* (with complement) and *in vivo* ^18,19^. To examine their potential as a target for antibody dependent cellular cytotoxicity (ADCC), we used FcγRIIIa receptor-expressing Jurkat reporter cells as a surrogate for ADCC effector cell recognition of anti-N mAb coated SARS-CoV-2 infected cells. Vero and BHK-21_hACE2 SARS-CoV-2 infected cells activated reporter cells in an anti-N mAb concentrationdependent manner (Fig. 4d). Activation was not observed in the absence of infection or with a control human mAb with the same heavy chain.

Taken together, these findings indicate that N protein from each of the highly pathogenic human CoVs blocks chemokine function. This is consistent with the possibility that cell surface N blocks chemokine function *in vivo*, facilitating viral replication and transmission. Conversely, we also show that cell surface N is a potential target for ADCC, which potentially contributes to limiting viral replication and transmission.

## DISCUSSION

Here, we show that N synthesized during SARS-CoV-2 infection or from a transfected cDNA is expressed on the surface of both N synthesizing cells and neighboring cells. N binding to the cell surface is based on specific association with heparan sulfate and heparin. The most parsimonious explanation for these findings is that N is released from cells and binds to both producing and bystander cells from liquid phase. Remarkably, based on the flow cytometry, levels of N on SARS-CoV-2 infected cells equals or exceeds cell surface S on all but one of the seven cell types examined. This, in part is due to the retention of a substantial fraction of S in the early secretory pathway but also reflects a robust amount of cell surface N, likely in the range of 10^4^-10^5^ copies per cell.

The mechanism underlying N secretion remains to be established. N has two potential sites for addition of N-linked oligosaccharides in the secretory pathway that are glycosylated and readily detected when N is targeted to the ER with an artificial N-terminal signal sequence^20^. Without a signal sequence, N is not glycosylated. This indicates that, as with other viral nucleic acid-binding proteins (*e.g*., SV40 T antigen, influenza virus N), SARS-CoV-2 N is likely secreted through a non-canonical secretory pathway, possibly one of the three defined pathways that bypass insertion into the ER^21^. Interestingly, like N, several proteins non-canonically exported to the cell surface (HIV Tat, FGF2, and tau) bind heparan sulfate, which has been shown to be involved in traversing the plasma membrane^22^. It will be interesting to examine the extent to which cell surface export of N and other viral RNA binding proteins, as well as their cell surface binding, is based on heparan sulfate association.

N is typically the most abundantly expressed SARS-CoV-2 protein, and its transfer to non-infected cells potentially amplifies its contributions to viral fitness. The remarkable ability of cell surface N to bind chemokines and block chemotaxis of immune effector cells offers an evolutionary explanation for its cellular export and binding to infected and neighboring uninfected cells. Like N, chemokines are immobilized on source cells and their neighbors by binding to GAGs. A number of viruses are known to express chemokine-binding proteins, which modulate chemokine activity by interacting with the GAG- or receptor-binding domain of chemokines, or both ^23,24^. Our findings establish SARS-CoV-2 N as the first CoV chemokine binding protein, one with a remarkably high affinity (nanomolar range) for multiple chemokines.

The binding of N to heparin, which limits coagulation at inflammation sites, suggests a possible role for secreted N in promoting COVID-associated clotting abnormalities. N is present in intestine and lungs from recovered and fatal COVID-19 patients, respectively, while virus-like particles are rarely detected^25,26^, consistent with this intriguing possibility as well as a role in the chronic low level inflammation that causes “long COVID-19” symptoms.

The remarkable efficacy of spike only-vaccines demonstrates that antibodies to N are not required for COVID-19 protection. SARS-CoV-2 induces a robust anti-N Ab response, in part likely due to cross-reaction with memory B cells induced by seasonal CoV infections. These Abs may reduce SARS-CoV-2 disease in naïve individuals since we establish N as a potential target for antibody-mediated effector functions, including complement and NK cell-mediated lysis of infected cells. Abs, therefore may play an unexpected role in protection to SARS-CoV-2 infection afforded by immunization with N-expressing vectors presumed to function via induction of N-specific T cells^27–30^. N is an attractive vaccine target due to its strong immunogenicity and much lower antigenic drift than spike. This may be particularly relevant given the remarkable capacity of SARS-CoV-2 to acquire amino changes in S as illustrated by the recent introduction of the omicron variant with over thirty non-synonymous mutations in S.

In summary, our findings demonstrate unexpected roles for N in innate and adaptive immunity to SARS-CoV-2 and other human CoVs that may contribute to both pathogenesis and protection, and support N as an Ab and T cell target for future “universal” vaccines that provide broad protection against both future strains of SARS-CoV-2 as well as other human CoVs.

## METHODS

### Cells

Vero cells (# CCL-81), BHK-21 (# CCL-10), Caco-2 (# HTB-37), Calu-3 (# HTB-55), CHO-K1 (# CCL-61), CHO-pgsA-745 (# CRL-2242), CHO-pgsB-618 (# CRL-2241), CHO-pgsD-677 (# CRL-2244), CHO-pgsE-606 (# CRL-2246), HEK293-FT (# CRL-11268), A549 (# CCL-185) and MOLT-4 (# CRL-1582) cells were from the American Type Culture Collection (ATCC). MonoMac-1 cells (# ACC 252) were from the DSMZ-German Collection of Microorganisms and Cell Cultures. PBMCs were obtained from a healthy donor with informed consent, at the Department of Transfusion Medicine (NIH). Vero, BHK-21, Caco-2, Calu-3 and HEK293-FT cells were grown in DMEM with GlutaMAX (Thermo Fisher # 10566016). CHO-K1, CHO-pgsA-745, CHO-pgsB-618, CHO-pgsD-677, CHO-pgsE-606 and A549 cells were grown in F-12K medium (Thermo Fisher # 21127022). PBMCs, MOLT-4 and MonoMac-1 cells were grown in RPMI 1640 (Thermo Fisher # 11875119). BHK-21_hACE2, CHO-K1_hACE2, HEK293-FT_hACE2 and A549_hACE2 cells were grown in their correspondent medium with 250-500 μg/ml of blasticidin (Invivogen # ant-bl-1). All cell media were supplemented with 8% (v/v) not heat inactivated FBS (Hyclone # SH30071.03), but Caco-2 with 20%, and cells were grown cultured at 37° C with 5% CO_2_ in sterile flasks. Cells were passaged at ~80-90% confluence and seeded as explained for each individual assays.

### SARS-CoV-2 stock preparation

SARS-CoV-2 (isolate USA-WA1/2020, # NR-52281), SARS-CoV-2_eGFP (# NR-54002) and the Alpha variant (B.1.1.7, # NR-54000) were obtained from BEI resources. The Beta (B.1.351) and Delta variants (B.1.617.2) were obtained from Andrew Pekosz (Johns Hopkins University, US). Viruses were propagated by the NIAID SARS-CoV-2 Virology Core Laboratory under BSL-3 conditions using Vero (CCL-81) or Vero overexpressing human TMPRSS2 cells, cultured in DMEM supplemented with GlutaMAX, 2% FBS, penicillin, streptomycin, and fungizone. Virus stocks were deep-sequenced and subjected to minor variants analysis by the NIAID SARS-CoV-2 Virology Core Laboratory. The TCID50 and PFU of virus in clarified culture medium was determined on Vero cells after staining with crystal violet. SARS-CoV-2 infections were performed in the NIAID SARS-CoV-2 Virology Core BSL3 laboratory strictly adhering to its standard operative procedures.

### Generating mutant cell lines constitutively expressing hACE2

The Sleeping Beauty transposase system was used for the generation of BHK-21_hACE2, CHO-K1_hACE2, HEK293-FT_hACE2 and A549_hACE2 cells as previously described ^31,32^. In brief, a semi-confluent 60 mm plate was seeded with each cell line and co-transfected with 0.5 μg of pCMV(CAT)T7-SB100 (Transposase vector, Addgene # 34879) and 5 μg of pSBbi-Bla hACE2 (Transposon vector), using TransIT-LT1 Transfection Reagent (Mirus Bio), following manufacturer instructions. After 24 h, cells were transferred to a T-75 flask and selected with 250-500 μg/ml of blasticidin for two weeks. The surface expression of hACE2 was confirmed by flow cytometry using anti-human ACE2 AlexaFluor 647-conjugated Ab (R&D Systems, # FAB9332R). The expression was further confirmed by immunoblot using hACE2 Ab (Cell Signaling Technology, # 4355). The open reading frame of hACE2 (kindly provided by Sonja Best from NIAID/NIH) was cloned into pSBbi-Bla vector (Addgene # 60526) as described ^31^.

### Antibodies

Previously published Ab VH and VL amino acid sequences against SARS-CoV-2 N (# N18^33^) and SARS-CoV-2 S (# H4^34^) were commercially synthesized, cloned into a human IgG1 vector backbone, produced and purified (Synbio Technologies). A 100 μg aliquot of anti-SARS-CoV-2 N human mAb was conjugated with Alexa Fluor^®^ 647 Lightning-Link^®^ Conjugation Kit (Abcam # ab269823), while 100 μg of anti-SARS-CoV-2 S were conjugated with Alexa Fluor^®^ 488 Lightning-Link^®^ Conjugation Kit (Abcam # ab236553). Each experiment was repeated with similar results using anti-SARS-CoV-2 N rabbit polyclonal Ab (GeneTex # GTX135357) and anti-SARS-CoV-2 S rabbit polyclonal Ab (ProSci # 3525). Goat anti-mouse IgG Alexa Fluor 488-conjugated (Thermo # A-11001) or 647 (# A-21235), goat anti-rabbit IgG Alexa Fluor 488 (# A-11008) or 647 (# A-21245), and Goat anti-human IgG Alexa Fluor 488 (# A-11013) or 647 (# A-21445) were used as a secondary Abs.

The SARS-CoV-2 S stabilized (S^st^) sequence^35^ (R710G, R711S, R713S, K1014P, V1015P) was commercially cloned into a mammalian expression vector, produced and purified (GenScript Biotech). Mouse polyclonal anti-SARS-CoV-2 S^st^ serum was produced as followed: 8-to-12-week C57B6 mice (Taconic Farms Inc) were immunized with 4 μg of S^st^ diluted in DPBS, adjuvanted by TiterMax^®^ Gold (MilliporeSigma # T2684) (2:1) in 50 μl volume via intramuscular injections. Serum was collected 21 d after booster immunization, heat inactivated at 56° C for 30 min, aliquoted and stored at 4° C. Abs and serum were titrated and specificity was tested, by flow cytometry on HEK293-FT cells transiently expressing SARS-CoV-2 N (Addgene # 141391), S (BEI # NR-52310) or S^st^.

### Immunofluorescence

For confocal microscopy imaging, 2.5 x 10^4^ cells were seeded on 12 mm glass coverslips in 24-well plates in indicated medium with gentamycin (25 μg/ml) overnight. Cell were infected with SARS-CoV-2 at an MOI of 1 PFU/cell for 1 h at 37° C. Virus was aspirated, and cells then incubated in cell growth media. After 24 h, the cells were washed with DPBS (Gibco # 14190-144) containing 5% goat serum (Jackson ImmunoResearch Labs. 005-000-121). Primary and secondary Abs were incubated with live cells at 4° C for 30 min. Cells were then washed twice with DPBS/5% goat serum and fixed in 4% PFA for 30 min at room temp. After fixation, coverslips were washed in DPBS and deionized _i_H_2_O, and mounted with Dapi Fluoromount G™ mounting medium (VWR # 102092-102). Images were acquired with a Leica STELLARIS 8 confocal microscope platform equipped with ultraviolet and white light lasers, using a 63x oil immersion objective (Leica Microsystems # 11513859), with a 1x zoom resolution of 512 x 512 pixels. Maximum intensity projections (MIPs) were processed from z-stacks (at least 15 0.3 μm z-steps per image); and for background correction (Gaussian filter) and color processing, using Imaris (Bitplane). Background levels of signal for each cell type were set based on mock-infected stained conditions. Animations (gifs) were generated with Photoshop 2022 (Adobe). Tile scans were taken of representative infected areas and individual fields (tiles) were merged into one image. Mock-infected coverslips were processed in parallel with infected counterparts, and SARS-CoV-2-infected coverslips were also incubated with all secondary Abs only as controls, and images were acquired using identical photomultiplier and laser settings.

### Flow cytometry

For cell surface protein expression analyses, 1 x 10^5^ cells were seeded on 24-well plates and mock or SARS-CoV-2-infected at an MOI of 1 pfu for 1 h at 37° C, followed by apsirating the virus inoculum and adding medium containing 2% FBS. After the indicated hpi, the cells were washed with DPBS, trypsinized with TrypLE™ Express Enzyme (Thermo Fisher # 12604039) or 0.25% Trypsin-EDTA (Thermo Fisher # 25200056) for 5 min at 37° C, transferred to a 96 well plate and washed with HBSS (Lonza # 10-527F) with 0.1% BSA. Cells were stained live with Alexa Fluor-conjugated Abs (or primary and secondary Abs), and LIVE/DEAD™ Fixable Violet Dead Cell Stain Kit (Thermo Fisher # L34964) in DPBS, for 25 min at 4° C. After Ab staining, cells were twice washed with HBSS 0.1% BSA and then fixed in 4%PFA for 30 min at room temp. PFA was aspirate and cells were resuspended in HBSS 0.1% BSA for analysis. To control for possible removal of cell surface antigens by trypsinization, in parallel infected cells after 24 h were washed with DPBS and directly stained in the 24 well plate prior to trypsinization. This resulted in similar levels of cell surface viral protein detection between trypsinized-stained and vice versa conditions.

For cell surface protein binding assays using recombinant proteins, indicated cells were trypsinized, washed with DPBS, and 1 x 10^5^ cells transferred to 96-well plates. Indicated amounts of recombinant GFP-His (Thermo Fisher # A42613) or SARS-CoV-2 N-His (Sino Biological # 40588-V08B, Acro Biosystems # NUN-C5227, Ray Biotech # 230-30164) were resuspended in 100 μl of DPBS, and cells were incubated for 15 min at 37° C and orbital shaking of 150 rpm. Cells were washed twice and stained as described above for subsequent flow cytometry analysis. For electric charge neutralization assays, after incubation with recombinant proteins and twice washed, cells were incubated with 10 μg/ml of polybrene (MilliporeSigma # TR-1003-G) in DPBS for 15 min at 37° C and orbital shaking of 150 rpm. Then, cells were washed twice and stained as described above for subsequent analysis.

For heparinase assays, 1 x 10^5^ single cells in 96-well plates were treated with Bacteroides heparinase I (4.8 units), II (1.6 units) and III (0.28 units) (NEB # P0735S, # P0736S, # P0737S) in DPBS for 1 h at 30° C. Cells were washed twice, incubated with recombinant proteins, and stained as described above for subsequent analysis.

For transient surface protein expression, 2 x 10^5^ cells were seeded on 12-well plates and transiently transfected with 2 μg of plasmids encoding SARS-CoV-2 N (Addgene # 141391) or eGFP (Addgene # 141395) with TransIT-LT1. At indicated time post transfection, cells were processed as described above for cell surface protein binding assays.

For every assay and condition, at least 30,000 cells were analyzed on an BD FACSCelesta™ Cell Analyzer (BD Biosciences) with a high throughput system unit, and quadrants in double staining plots were set based on mock-infected condition for each cell type. Data were analyzed with FlowJo (Tree Star) and plotted with Prism v9.1.1 software (GraphPad).

### Cell co-culture assays

For infectable and non-infectable cell co-culture assays, 9 x 10^5^ cells of each indicated SARS-CoV-2-non-infectable cell type were stained with CellTrace™ Violet (Thermo Fisher # C34557), following manufacturer’s instructions, and seeded in 6-well plates. Then, 1 x 10^5^ CHO-K1_hACE2 cells (SARS-CoV-2-infectable) were homogeneously mixed and co-seeded with indicated non-infectable cell type, being co-cultured overnight. Co-cultured cells were inoculated with SARS-CoV-2 at an MOI of 1 for 1 h at 37° C, followed by removal of the virus inoculum and replacement of the medium containing 2% FBS. After 24 h, cells were washed with DPBS and directly stained live on their 6 wells with Alexa Fluor^®^-conjugated Abs and LIVE/DEAD™ Fixable Violet Dead Cell Stain Kit, in DPBS for 25 min at 4° C. After staining, cells were washed twice with HBSS 0.1% BSA, trypsinized with TrypLE™ Express Enzyme for 5 min at 37° C, transferred to 96 wells, washed with HBSS 0.1% BSA and fixed in 4% PFA for 30 min at room temp. PFA was aspirated and cells were resuspended in HBSS 0.1% BSA for flow cytometry analysis.

For transfected and non-transfected cell co-culture assays, 2 x 10^5^ cells were seeded on 6-well plates and transiently transfected with 2 μg of plasmids encoding SARS-CoV-2 N or eGFP with TransIT-LT1. After 24 h, 9 x 10^5^ non-transfected cells were stained with CellTrace™ Violet and co-seeded with their transfected homologs, being co-cultured for 24 h. Then, cells were washed with DPBS and directly stained live on their 6 wells with Alexa Fluor^®^-conjugated Abs and LIVE/DEAD™ Fixable Violet Dead Cell Stain Kit, in DPBS for 25 min at 4° C. Cells were washed twice with HBSS 0.1% BSA, trypsinized with TrypLE™ Express Enzyme for 5 min at 37° C, transferred to 96 wells, washed twice with HBSS 0.1% BSA and resuspended in HBSS 0.1% BSA for flow cytometry analysis.

For every assay and condition, at least 100,000 cells were analyzed on an BD FACSCelesta™ Cell Analyzer with a high throughput system unit, and data were analyzed with FlowJo and plotted with GraphPad Prism software.

### Cytokines and GAGs

Recombinant human cytokines used in this study (CCL1, CCL2, CCL3, CCL3L1, CCL4, CCL4L1, CCL5, CCL7, CCL8, CCL11, CCL13, CCL14, CCL15, CCL16, CCL17, CCL18, CCL19, CCL20, CCL21, CCL22, CCL23, CCL24, CCL25, CCL26, CCL27, CCL28, CXCL1, CXCL2, CXCL3, CXCL4, CXCL5, CXCL6, CXCL7, CXCL8, CXCL9, CXCL10, CXCL11, CXCL12α, CXCL12β, CXCL13, CXCL14, CXCL16, XCL1, CX3CL1, IL-1α, IL-1β, IL-6, IL-6Rα, IL-10, IL-12p70, IL-13, IL-17a, IL-18BP-Fc, IL-23, IL-27, IL-35, TNF-α, TNF-β, IFN-β, IFN-γ, IFN-λ1, IFN-ω) from PeproTech, and (IFN-α2 and IL-18) from Sino Biological, were reconstituted in DPBS 0.1% BSA at 10 μM, aliquoted and stored at −80° C. Heparin (# 2106), heparan sulfate from bovine kidney (# H7640), chondroitin sulphate A (# C9819) and chondroitin sulphate B (# C3788) were obtained from MilliporeSigma. Heparan sulfate from porcine mucosa (# AMS.GAG-HS01) and keratan sulfate (# AMS.CSR-NAKPS2-SHC-1) were purchased from ASMBIO. We assumed an average molecular weight of 30 kDa for heparan sulfate from porcine mucosa and 15 kDa for heparin ^36^.

### BLI assays

High throughput binding assays were performed on an Octet Red384 (ForteBio) instrument at 30° C with shaking at 1,000 rpm. Streptavidin (SA) biosensors (Sartorius # 18-5019) were hydrated for 15 min in kinetics buffer (DPBS, 1% BSA, 0.05% Tween-20). SARS-CoV-2 structural proteins and accessory factors (2X-StrepTag tagged) in lysis buffer from commercial sources or crude lysates of transfected cells (see details below) were loaded into SA biosensors up to 0.5-5 nm of binding response for 300-600 s, prior to baseline equilibration for 180 s in kinetics buffer. Association of each analyte in kinetics buffer at indicated concentration was carried out for 300 s, followed by dissociation for 300 s or longer. Standard binding and kinetic assays between SARS-CoV-2 N and GAGs or chemokines were performed as described above for binding assays. Negative signal of N binding to GAGs, expected given the large size of heparin molecules, was flipped prior further analysis ^37^. The data were baseline subtracted prior to fitting performed using the homogeneous (1:1) and heterogeneous binding models (2:1, mass transport, 1:2) within the ForteBio Data Analysis HT software v12.0.1.55. Mean K_D_, k_on_, k_off_ values were determined with a global fit applied to all data. The performance of each binding model fitting to the data was assessed based on the lowest sum of the squared deviations or measure of error between the experimental data and the fitted line (χ^2^), and the highest correlation between fit and experimental data (R^2^).

The experiments were repeated with at least three independently produced batches of recombinant protein in crude lysates, obtained from 30 x 10^6^ HEK293-FT cells transfected with 30 μg of plasmids encoding SARS-CoV-2 structural proteins and accessory factors with TransIT-LT1. SARS-CoV-2 St containing 2X-StrepTag at the C-terminal region was commercially synthesized as mentioned above (GenScript Biotech). SARS-CoV-2 N, M, E, ORF3a, ORF3b, ORF6, ORF7a, ORF7b, ORF8, ORF9b, ORF9c and ORF10 plasmids without signal peptide for secretion, described in ^38^, were acquired from Addgene (www.addgene.org/Nevan_Krogan). After 24 h, transfected cells were selected with 10 μg/ml of puromycin (Invivogen # ant-pr-1). After 48 h, transfected cells were trypsinized, washed with DPBS and lysated for 30 min at 4° C in 1 ml of lysis buffer (50 mM Tris-HCl pH 7.4, 150 mM NaCl, 5 mM KCl, 5 mM MgCl_2_, 1% NP-40 and 1x protease inhibitors (Roche # 4693159001)), followed by centrifugation at 1000 x *g* at 4° C. Clarified supernatants (crude lysates) were collected, aliquoted, stored at −20° C and characterized by immnoblotting using mouse anti-2xStrep tag (Qiagen #34850, 1:1,000), and secondary goat anti-mouse IgG IRDye® 800CW (LI-COR # 926-32210, 1:10,000). rN was additionally characterized by immunoblotting using IRDye® 680RD Streptavidin (LI-COR # 926-68079), and human anti-SARS-CoV-2 N mAb (N18) followed by IRDye® 680RD Goat anti-Human IgG Secondary Ab (LI-COR # 926-68078).

For GAGs competition assays of chemokine binding, selected chemokines (100 nM) were incubated in kinetics buffer alone or with indicated concentrations of soluble heparan sulfate from bovine kidney, chondroitin sulfate A and chondroitin sulfate B for 10 min at room temp. The mixture was used to measure the association of N (in nM of binding response), as described above for the BLS binding assays. The value in nm of binding response of each chemokine binding without GAGs was considered 100%.

### Chemotaxis assays

Recombinant human CXCL12β (3 nM), alone or in combination with purified recombinant proteins were placed in the lower chamber of a 96-well ChemoTx System plate (Neuro Probe # 101-3 and # 101-5) in RPMI 1640 1% FBS. As internal controls within each assay, medium or recombinant protein alone were used. PBMCs, MonoMac-1 and MOLT-4 cells (1.25 x 10^5^) were placed on the upper compartment and separated from the lower chamber by a 3 or 5 μm pore size filter. The cells were incubated at 37° C for 3 h in a humidified incubator with 5% CO_2_. The migrated cells in the lower chamber were stained with 5 μl of CellTiter 96 AQueous One Solution Cell Proliferation Assay (Promega # G3580) for 2 h at 37° C with 5% CO_2_, measuring absorbance at 490 nm using a Synergy H1 plate reader (Bio-Tek).

The following recombinant proteins were used: SARS-CoV-2 S1-His (Sino Biological # 40591-V08B1), SARS-CoV-2 N-His (Sino Biological # 40588-V08B), SARS-CoV-2 N-His (Acro Biosystems # NUN-C5227), SARS-CoV-2 N-His (Ray Biotech # 230-30164), SARS-CoV-1 N-His (Acro Biosystems # NUN-S5229) and MERS-CoV N-His (Acro Biosystems # NUN-M52H5). SARS-CoV-2 N-His from Sino Biological (# 40588-V08B) was used in all assays unless indicated.

### ADCC reporter assay

For each indicated cell type, 1 x 10^4^ cells were seeded on 96-well flat white tissue culture-treated plates (Thermo Fisher # 136101), cultured overnight, and infected with SARS-CoV-2 at an MOI of 1 (target cells). At 24 hpi, infected target cells were washed with DPBS and the medium was replaced with 50 μl of RPMI 1640 with 4% low IgG serum (Promega # G711A) containing 5 x 10^4^ Jurkat effector cells (Promega # G701A) and serial dilutions of indicated human mAbs. After overnight incubation at 37° C with 5% CO_2_, 50 μl of Bright-Glo™ Luciferase Assay lysis/substrate buffer (Promega # E2620) were added and luminescence was measured after 10 min using a POLARstar Omega plate reader (BMG LABTECH) within the luciferase Glow assay template and the following parameters: gain 3600; measurement interval time 0.1 s; and maximum counts 2 x 10^6^. Measurements were performed in triplicate and relative luciferase units (RLU) were plotted and analyzed with GraphPad Prism software. Data fitting with GraphPad Prism was performed with the non-linear regression dose-response-stimulation model.

### Statistical analysis

Statistical analyses were performed using GraphPad Prism software. When indicated, *p* values were calculated using Student’s two-tailed unpaired *t* test (at 99% confidence interval) and *p* < 0.01 was considered statistically significant. On the other hand, One-way ANOVA and Dunnett’s Multiple comparison test (at 95% confidence interval) was used to compare all conditions against untreated or mock-infected cells (as indicated for each case), considering *p* < 0.05 as statistically significant.

## Supporting information

Supplementary Animations

## ACKNOWLEDGMENTS

We thank James S. Gibbs for outstanding technical assistance. We are grateful to the NIAID SARS-CoV-2 Virology Core BSL3 Laboratory staff for their training, help, and support. This work was supported by the Division of Intramural Research of the National Institute of Allergy and Infectious Diseases.

## AUTHOR CONTRIBUTIONS

A.D.L.M. conceived, designed, and performed experiments, analyzed data, interpreted results, and wrote the manuscript.

I.K. produced in-house antibodies, and helped to design and analyze ADCC assays.

J.H. generated cell lines expressing hACE2.

J.W.Y. conceived experiments and wrote the manuscript.

## COMPETING INTERESTS

The authors declare no competing interests.

**Supplementary Information is available for this paper.**

## MATERIALS & CORRESPONDENCE

Correspondence and material requests should be addressed to J.W.Y.

---

**Extended Data Fig. 1:**
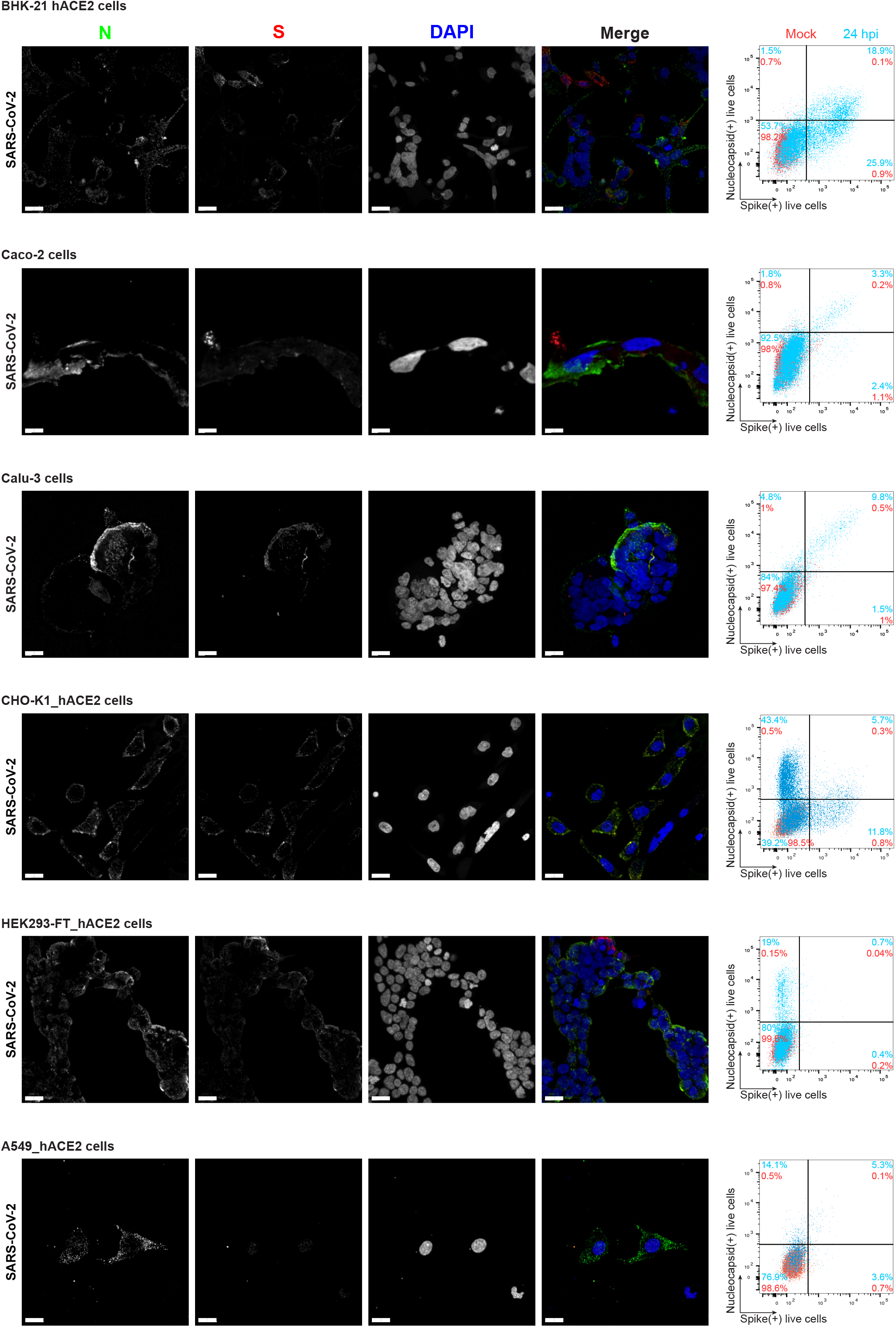
SARS-CoV-2 N protein is expressed on the cell surface of diverse infected cell types. Maximum intensity projections (MIP) of laser confocal microscopy z-stack images and histogram overlays of flow cytometry analyses of live wild-type SARS-CoV-2-infected BHK-21_hACE2, Caco-2, Calu-3, CHO-K1_hACE2, HEK293-FT_hACE2 and A549_hACE2 cells stained live with Abs at 24 hpi (MOI = 1). Scale bars = 20 μm. Images are representative of three independent experiments with similar results. Representative plots of flow cytometry analyses show double staining of surface S and N, indicating the percentage of the gated cell population for each quadrant of the double staining. Data are representative of one experiment out of at least three independent experiments performed in triplicate.

**Extended Data Fig. 2:**
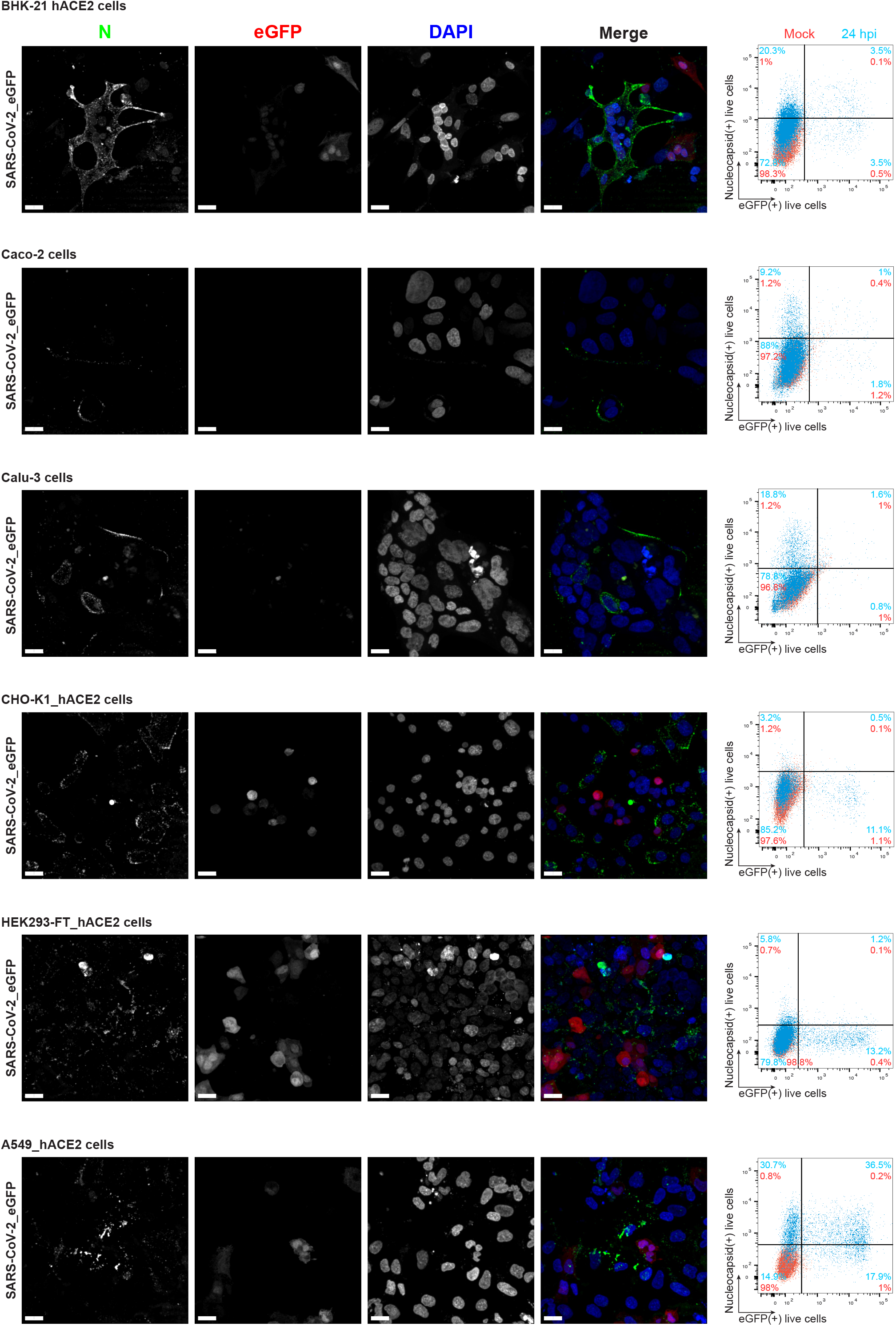
SARS-CoV-2 N protein is a robust marker of cell-surface infection across cell types. MIP of laser confocal microscopy z-stack images and histogram overlays of flow cytometry analyses of live eGFP expressing SARS-CoV-2-infected BHK-21_hACE2, Caco-2, Calu-3, CHO-K1_hACE2, HEK293-FT_hACE2 and A549_hACE2 cells stained live with MAb against N at 24 hpi (MOI = 1). Scale bars = 20 μm. Images are representative of two independent experiments with similar results. Representative plots of flow cytometry analyses show double staining of eGFP and surface N, indicating the percentage of the gated cell population for each quadrant of the double staining. Data are representative of one experiment out of at least three independent experiments performed in triplicate.

**Extended Data Fig. 3:**
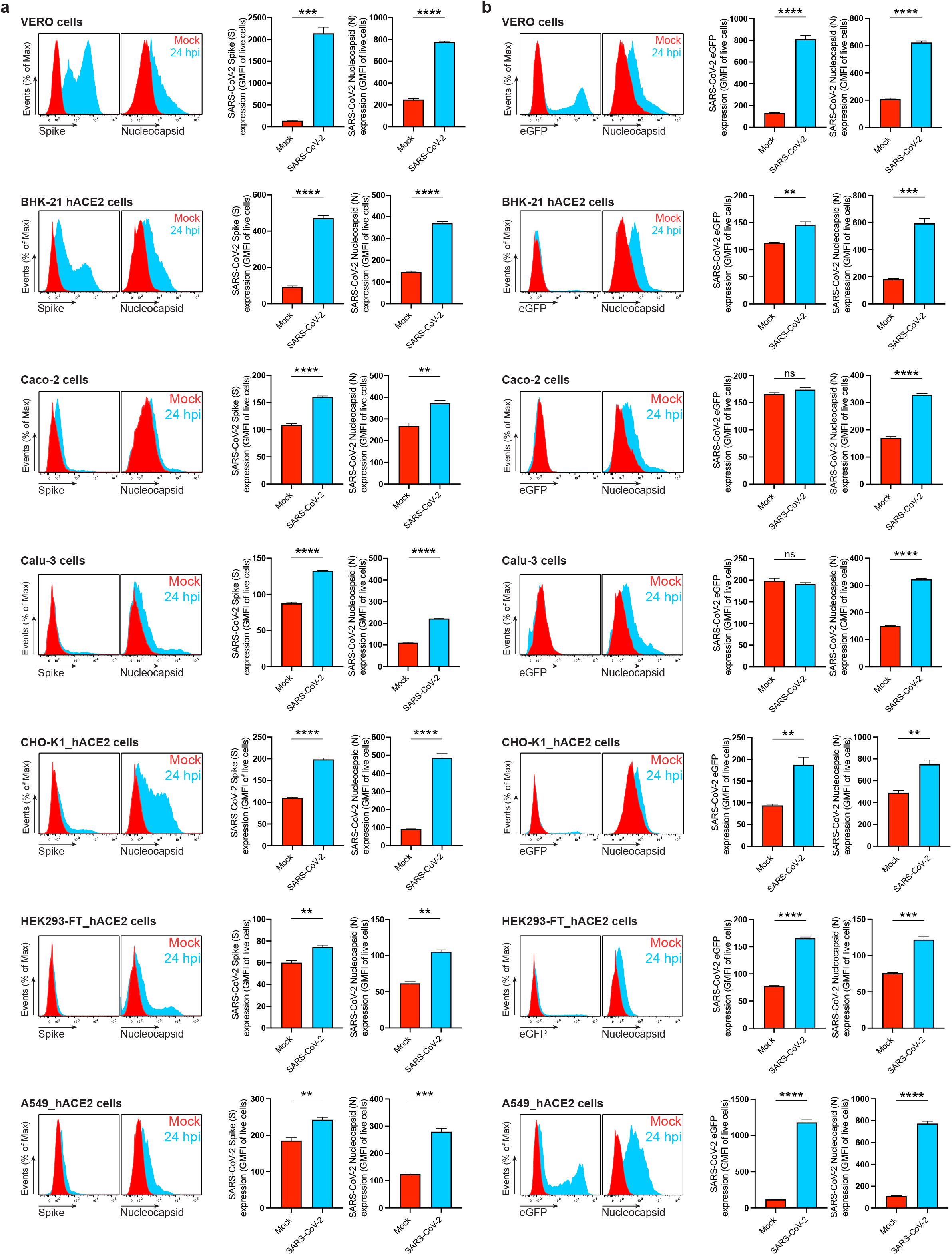
SARS-CoV-2 N is significantly detected on the cell surface of all live infected cell types tested in this study. Flow cytometry analyses of Vero, BHK-21_hACE2, Caco-2, Calu-3, CHO-K1_hACE2, HEK293-FT_hACE2 and A549_hACE2 cells inoculated with wild-type (**a**) or eGFP reported (**b**) SARS-CoV-2 (MOI = 1), and stained live with Abs at 24 hpi against SARS-CoV-2 S and N proteins. For each cell type and infection, the following is shown: histogram overlays of surface S and N, and intracellular eGFP, as well as the GMFI is plotted showing mean +/- SEM (n = 3). Histogram overlays correspond to **Fig. 1b** (Vero cells), while those shown in (**a**) to **Extended Data Fig. 1** and those in (**b**) to **Extended Data Fig. 2.** Data are representative of one experiment out of at least three independent experiments performed in triplicate. *ns* (nonsignificant) *p* > 0.01, ** *p* < 0.01, *** *p* < 0.001, **** *p* < 0.0001 by Student’s two-tailed unpaired *t*-test.

**Extended Data Fig. 4:**
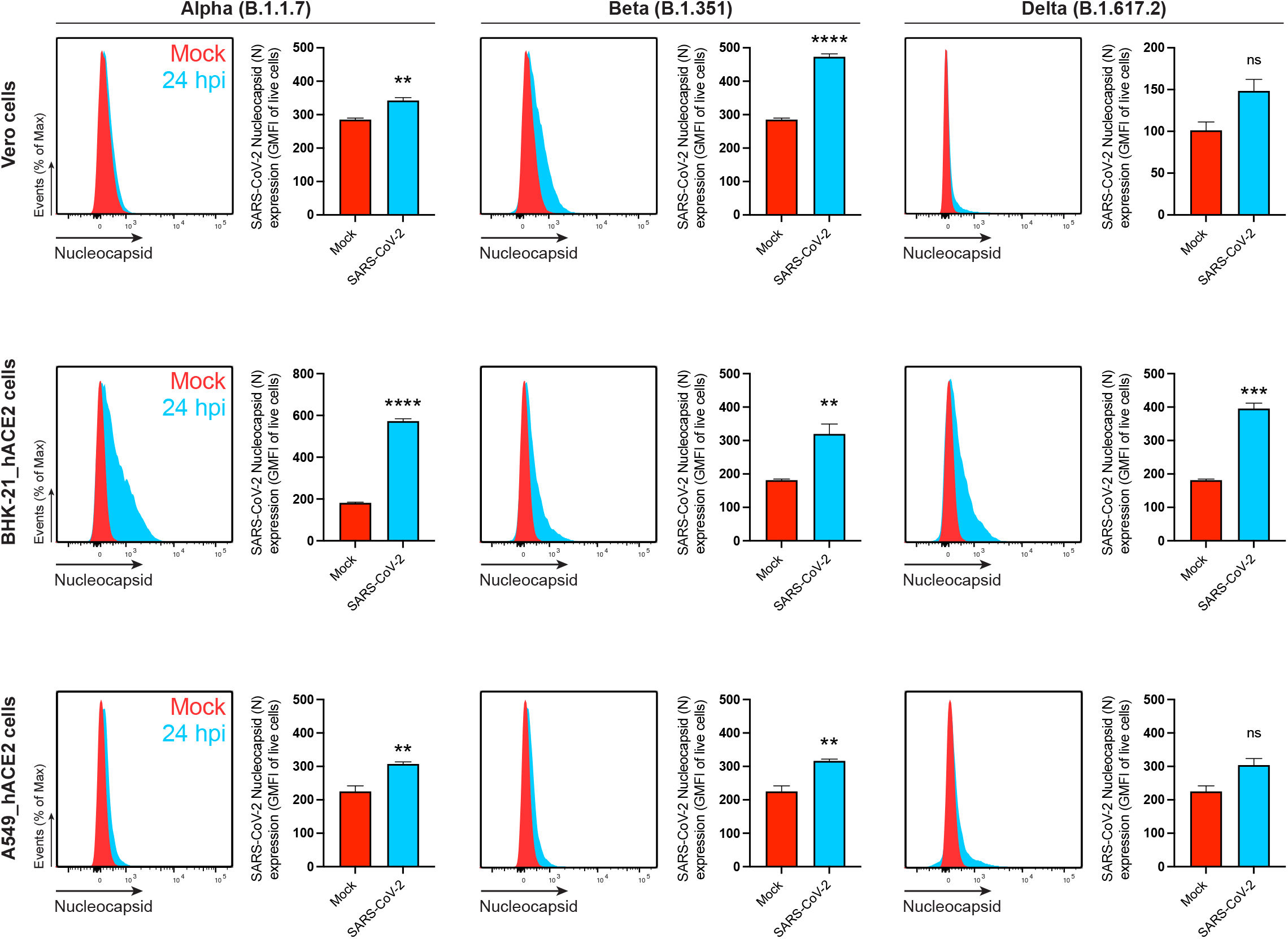
SARS-CoV-2 N is also detected on the cell surface of live cells infected with the Alpha, Beta and Delta variants. Flow cytometry analyses of Vero, BHK-21_hACE2, and A549_hACE2 cells inoculated with SARS-CoV-2 variants (MOI = 1), and stained live with Abs at 24 hpi against SARS-CoV-2 N. For each cell type and infection, the following is shown: histogram overlays of surface N on live cells, as well as the GMFI is plotted showing mean +/- SEM (n = 3). Data are representative of one experiment out of at least three independent experiments performed in triplicate. *ns* (nonsignificant) *p* > 0.01, ** *p* < 0.01, *** *p* < 0.001, **** *p* < 0.0001 by Student’s two-tailed unpaired *t*-test.

**Extended Data Fig. 5:**
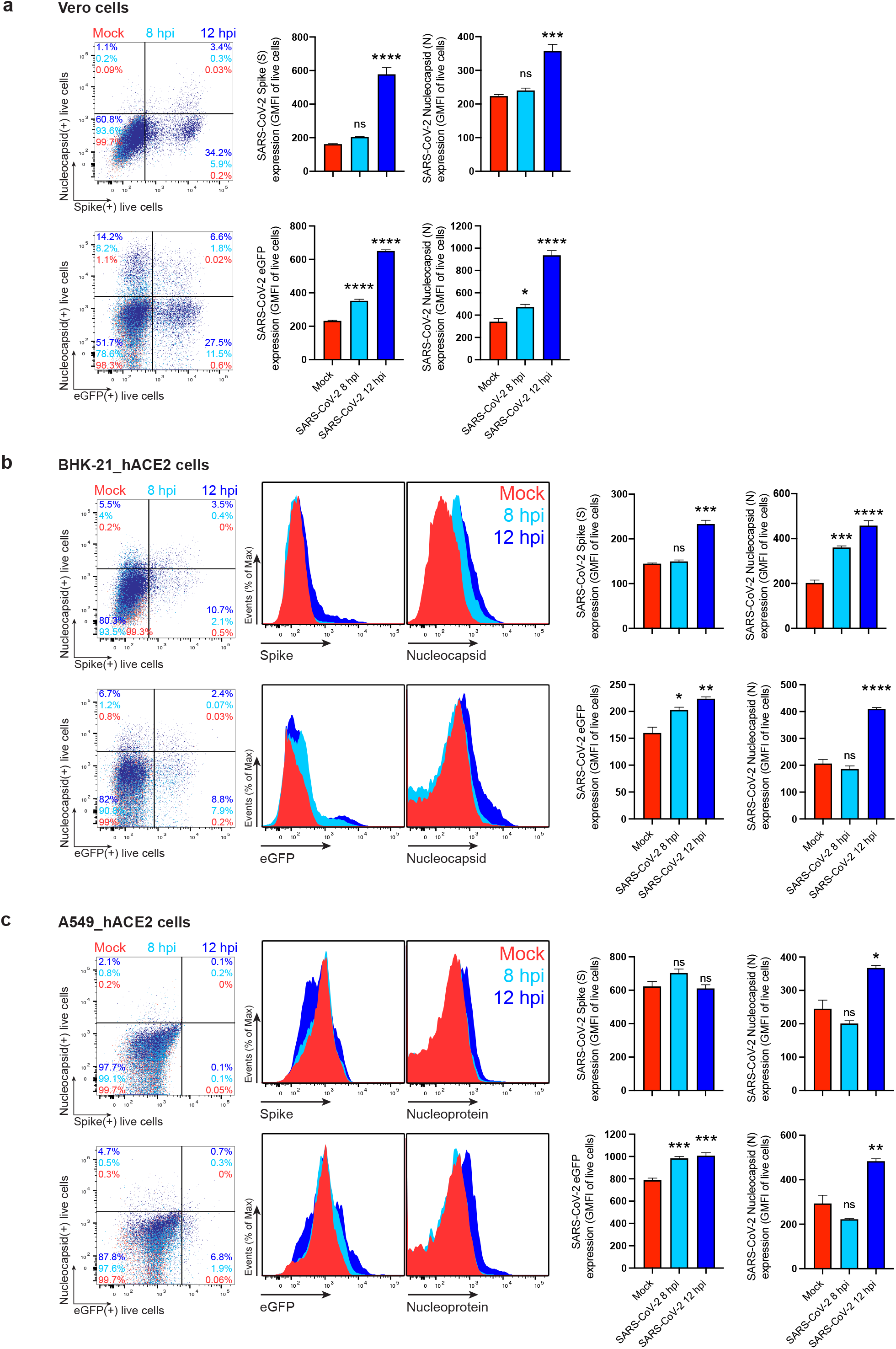
SARS-CoV-2 N reaches the cell surface as soon as 8 to 12 hpi. Time course of surface S, N, and eGFP proteins expression in live Vero (**a**), BHK-21_hACE2 (**b**) and A549_hACE2 cells (**c**) infected with wild-type and eGFP reporter SARS-CoV-2 at 8 and 12 hpi (MOI = 1). Representative plots of flow cytometry analyses show double staining of surface S, N, and intracellular eGFP, indicating the percentage of the gated cell population for each quadrant. Histogram overlays of surface S and N, and intracellular eGFP are shown, as well as the GMFI is plotted showing mean +/- SEM (n = 3). Histogram overlays from Vero cells analyses are shown in **Fig. 1a, b.** Data are representative of one experiment out of at least three independent experiments performed in triplicate. One-way ANOVA and Dunnett’s Multiple comparison test were used to compare all conditions against mock-infected cells: *ns* (nonsignificant) *p* > 0.05, * *p* < 0.05, ** *p* < 0.01, *** *p* < 0.001, **** *p* < 0.0001.

**Extended Data Fig. 6:**
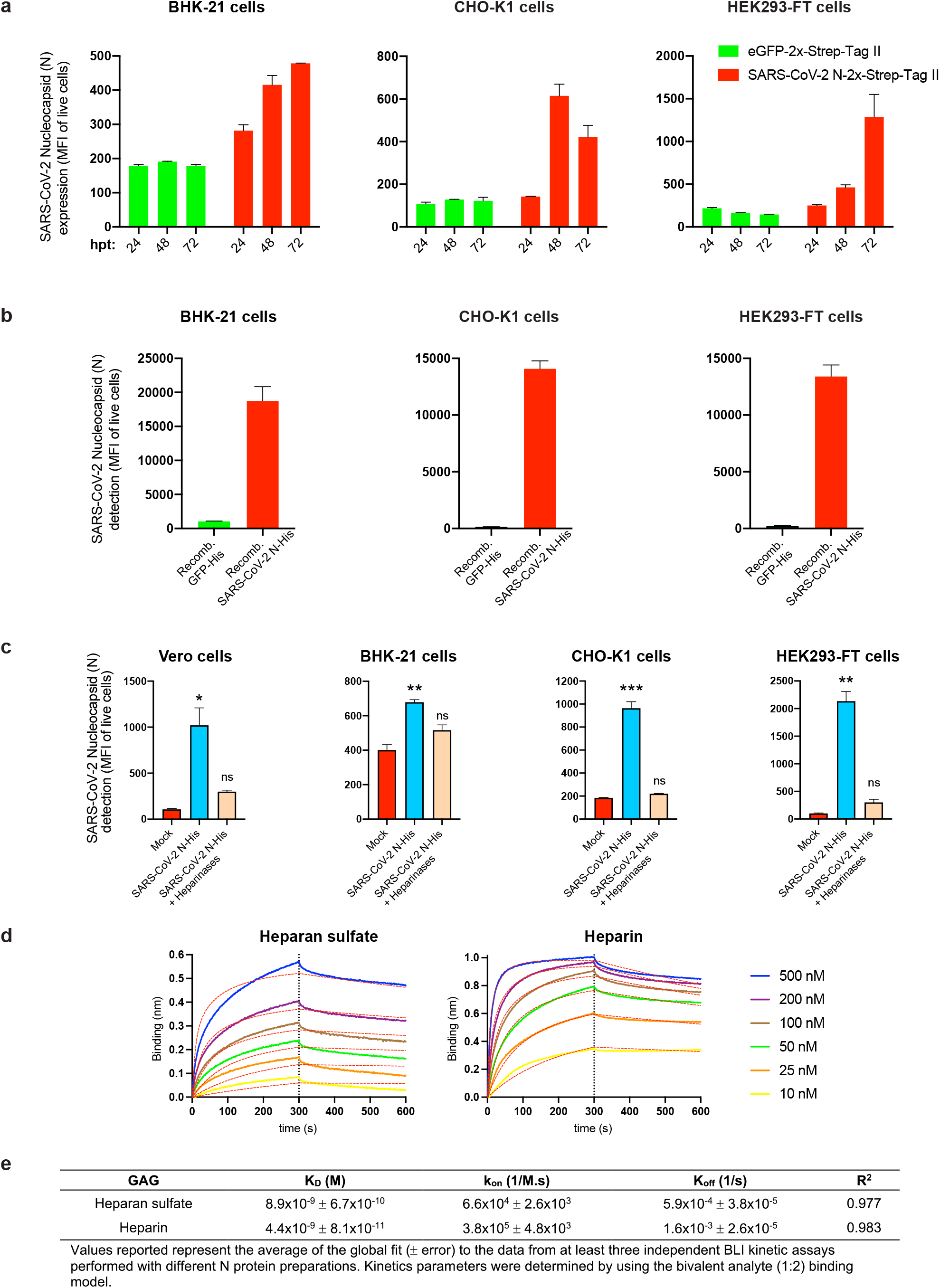
Other SARS-CoV-2 genes are not required for N cell surface expression and binding to surface heparan sulfate/heparin. **a,** Flow cytometry analyses of surface N kinetics (24-72 h) expression in BHK-21, CHO-K1 and HEK293-FT cells transiently transfected with a plasmid encoding eGFP or N protein, detected with Abs. **b,** Flow cytometry analyses of exogenous rN binding to BHK-21, CHO-K1 and HEK293-FT cells, incubated with recombinant eGFP or N protein (100 ng) for 15 min, washed twice, stained live with Abs. **c,** Flow cytometry analyses of BHK-21, CHO-K1 and HEK293-FT cells treated with heparinases for 1 h, washed twice, incubated with 50 ng of rN protein for 15 min, washed twice again, stained live with Abs, and analyzed. The MFI of expressed (**a**) or bound (**b, c**) surface N protein from live cells is plotted for each case, showing mean +/- SEM (n = 2). In (**c**), One-way ANOVA and Dunnett’s Multiple comparison test were used to compare all conditions against untreated cells (mock): *ns* (nonsignificant) *p* > 0.05, * *p* < 0.05, ** *p* < 0.01, *** *p* < 0.001. **d,** BLI sensorgrams of kinetic assays depicting the interaction between immobilized N protein and different concentrations of heparan sulfate and heparin. All curves were analyzed with the ForteBio Data Analysis HT software, where red dashed lines correspond to a global fit of the data using the bivalent analyte model (1:2). **e,** Table showing averaged values from the kinetic analyses of the N protein binding to heparan sulfate and heparin by BLI. All analyses were repeated with different protein preparations, and one representative assay out of at least three independent assays performed in duplicate is shown.

**Extended Data Fig. 7:**
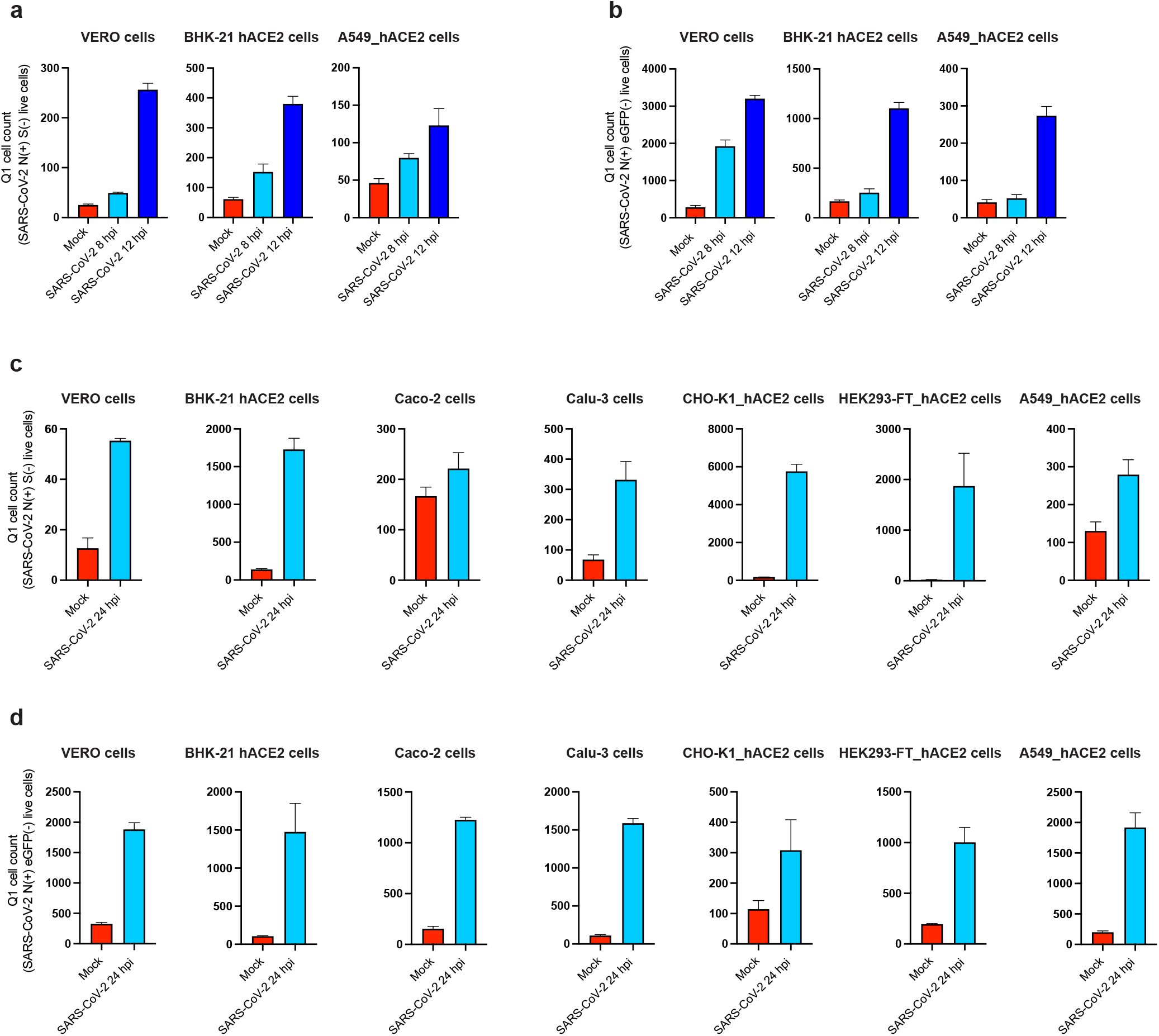
Additional evidence supporting transfer of N from infected to uninfected infected cells. **a – d,** Cells expressing N but no other marker of infection (S or eGFP) increase in a time-dependent manner. Quadrant 1 (Q1) in plots of flow cytometry analyses showing double staining of surface N and S/eGFP identifies a cell subset of cells only expressing N during infection. Bar histograms show mean +/- SEM (n = 3) from Q1 cell counts of cells at 8 and 12 h after infection with wild-type (**a**) or eGFP reporter virus (**b**), as well as at 24 hpi for each cell type analyzed after wild-type (**c**) or eGFP reporter virus (**d**) infection. Data are representative of one experiment out of at least three independent experiments performed in triplicate.

**Extended Data Fig. 8:**
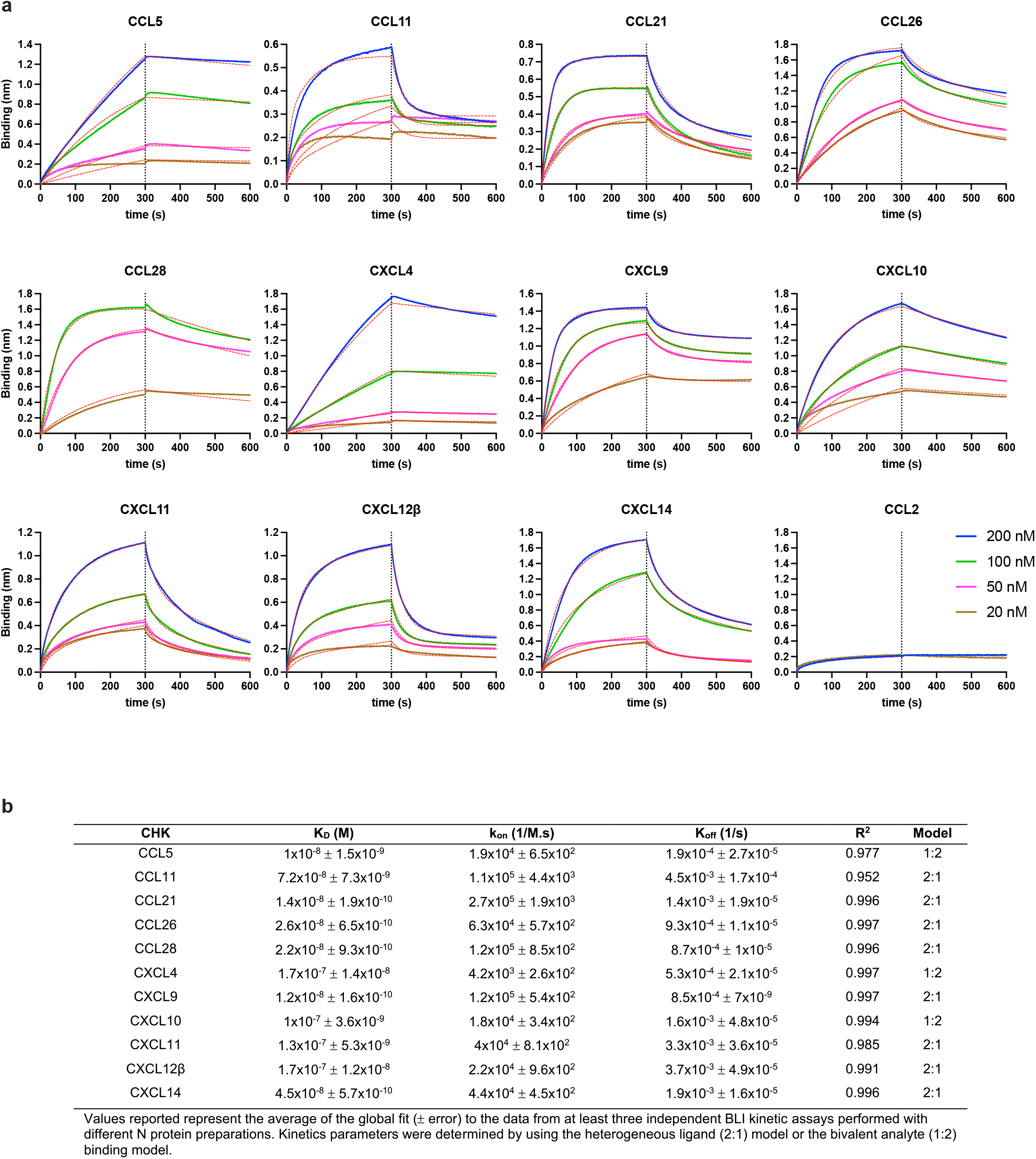
BLI demonstration that SARS-CoV-2 N protein binds to human chemokines with high affinity. **a,** BLI sensorgrams of affinity kinetic assays between immobilized SARS-CoV-2 rN protein and positively bound human chemokines identified from BLI HTS binding assays. Sensorgrams show association and dissociation phases. The vertical dotted line indicates the end of the association step. Curves were analyzed with the ForteBio Data Analysis HT software, where red dashed lines correspond to a global fit of the data using the heterogeneous ligand (2:1) or bivalent analyte binding model (1:2). CCL2 is shown as example of negative interaction. All analyses were repeated with different protein preparations, and one representative assay out of at least three independent assays performed in duplicate is shown. **b,** Table showing averaged values from the kinetic analyses of the N protein binding to each chemokine by BLI.

**Extended Data Fig. 9:**
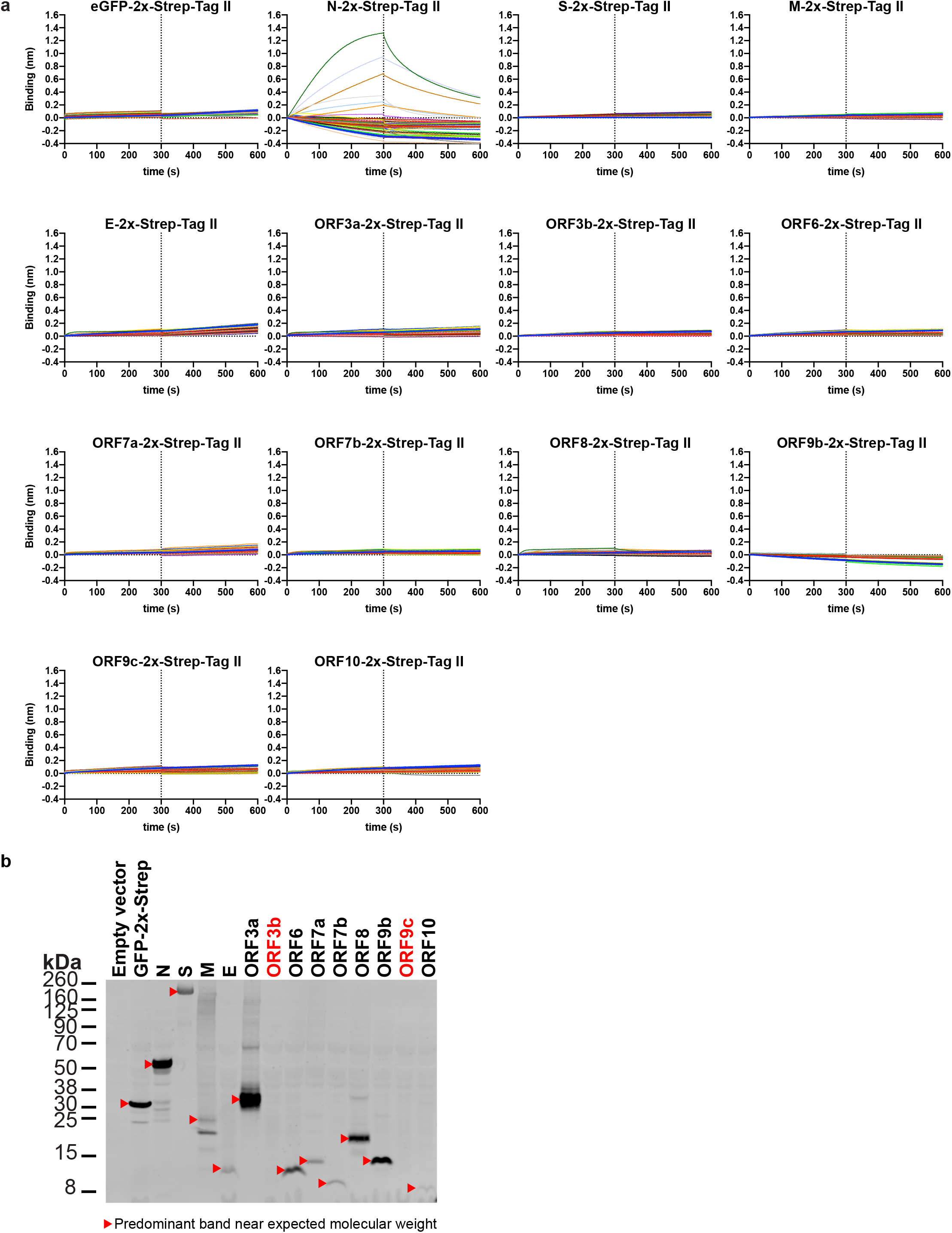
Among SARS-CoV-2 structural proteins and accessory factors SARS-CoV-2 N protein uniquely binds chemokines. **a,** BLI sensorgrams of HTS binding assays between immobilized eGFP, and SARS-CoV-2 structural proteins and accessory factors against 64 human cytokines at 100 nM (see detailed list in Material and Methods). N protein bound CCL5, CCL11, CCL21, CCL26, CCL28, CXCL4, CXCL9, CXCL10, CXCL11, CXCL12β and CXCL14 across different independent assays. Sensorgrams show association and dissociation phases. The dotted line indicates the end of the association step. The analyses were repeated with different protein preparations and one representative assay of two independent HTS is shown. **b,** Immunoblot detection of 2xStrep tag verified expression of predicted protein sizes (red arrowheads), which were loaded into streptavidin-coated biosensors for BLI HTS binding assays. Despite the lack of detection of ORF3b and ORF9c (labeled in red) by immunoblot, the expression of these ORFs was detected after positive loading into streptavidin-coated biosensors by BLI. For gel source data, see **Supplementary Fig. 3.**

**Extended Data Fig. 10:**
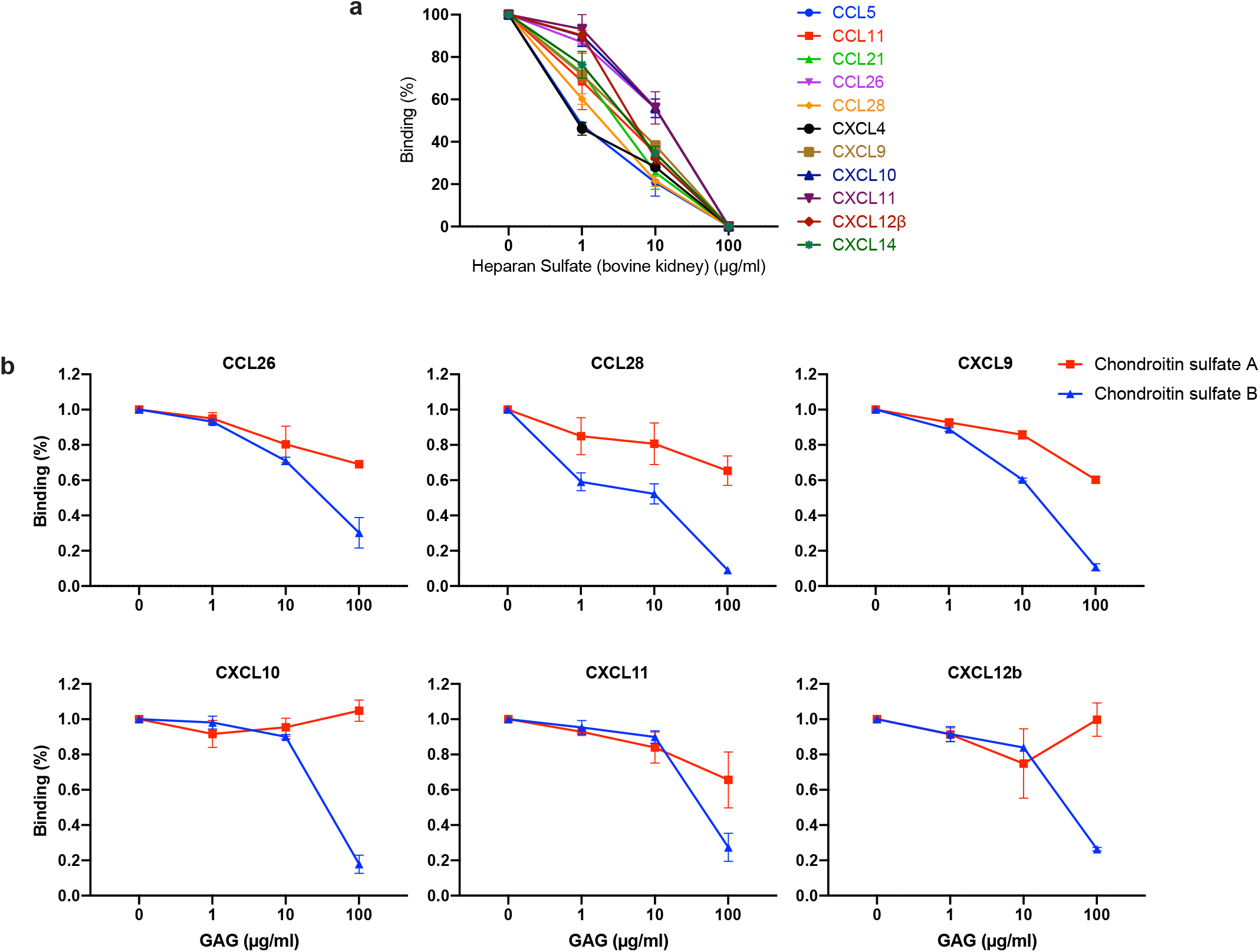
SARS-CoV-2 N protein binds chemokines through the GAG-binding domain of chemokines. Sulfated GAG competition of chemokine binding to N protein. chemokines at a concentration of 100 nM, alone or incubated with the indicated increasing concentrations of heparan sulfate from bovine kidney (**a**) or chondroitin sulfate A/B (**b**), were used for BLI binding analyses to N protein. The value of each chemokine binding without GAGs was considered 100%. Data represent the mean +/- SEM of 2-3 independent experiments. Heparan sulfate (from bovine kidney) interaction with N is considered negligible based on results from BLI assays shown in **Fig. 2f**.

