## Supplementary information for "Cell Surface SARS-CoV-2 Nucleocapsid Protein Modulates Innate and Adaptive Immunity"

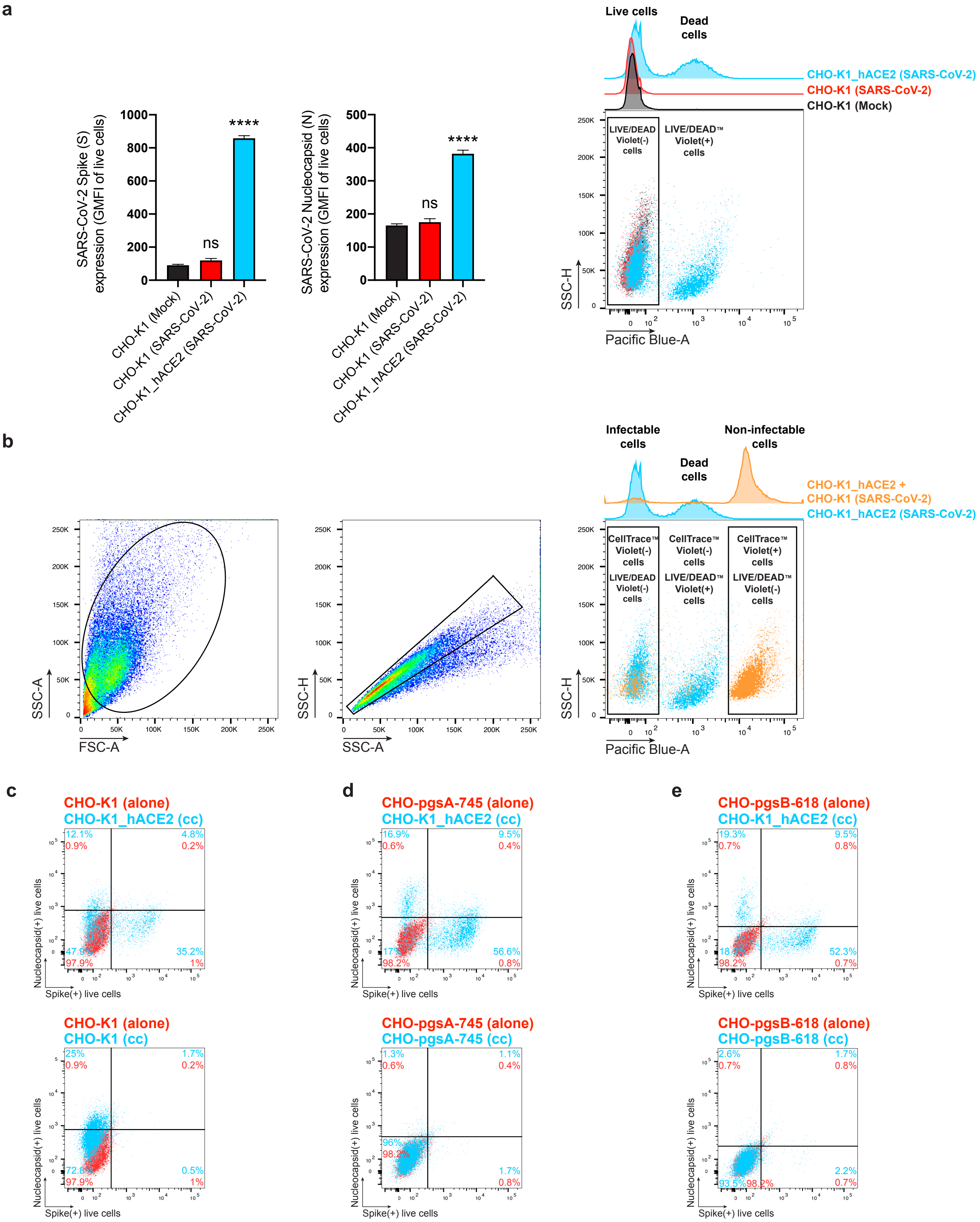


**Supplementary Fig. 1. Controls, gating strategies, and additional data for N transfer assays between SARS-CoV-2 infectable and non-infectable co-cultured cells. a,** Flow cytometry analyses of CHO-K1 and CHO-K1_hACE2 cells mock or SARS-CoV-2-infected (MOI = 1) and stained live at 24 hpi with Abs against SARS-CoV-2 S and N proteins and with LIVE/DEADTM Violet. The GMFI of live cells expressing S and N proteins is plotted, showing mean +/- SEM (n = 3). One-way ANOVA and Dunnett's Multiple comparison test were used to compare all conditions against mock-infected cells: *ns* (nonsignificant) $p > 0.05$, **** $p < 0.0001$. Representative plots and histogram semi-overlays of LIVE/DEAD Violet staining are shown, indicating the gating for live cells. One representative experiment of at least two independent experiments performed in triplicate is shown. **b,** Gating and staining strategy for live/dead, and infectable versus non-infectable cell exclusion for flow cytometry analyses shown in **Fig. 3**. CHO-K1, CHO-pgsA-745 and CHO-pgsB-618 cells were stained with CellTrace Violet prior co-culture with CHO-K1_hACE2 cells. Co-cultured cells were inoculated with SARS-CoV-2 (MOI = 1) and stained live at 24 hpi with Abs against SARS-CoV-2 S and N proteins and with LIVE/DEAD Violet, and analyzed by flow cytometry. Representative plots and histogram semi-overlays of LIVE/DEAD Violet and CellTraceTM Violet staining are shown, indicating the gating for live CellTrace Violet(-) cells (left), infected dead CellTrace Violet(-) cells (center), and uninfected live CellTrace Violet(+) cells cells (right). One representative experiment of two independent experiments performed in triplicate is shown. **c–e,** Representative plots of flow cytometry analyses showing double staining of surface S and N proteins from the same histogram overlays shown in **Fig. 3,** indicating the percentage of the gated cell population for each quadrant of the double staining.

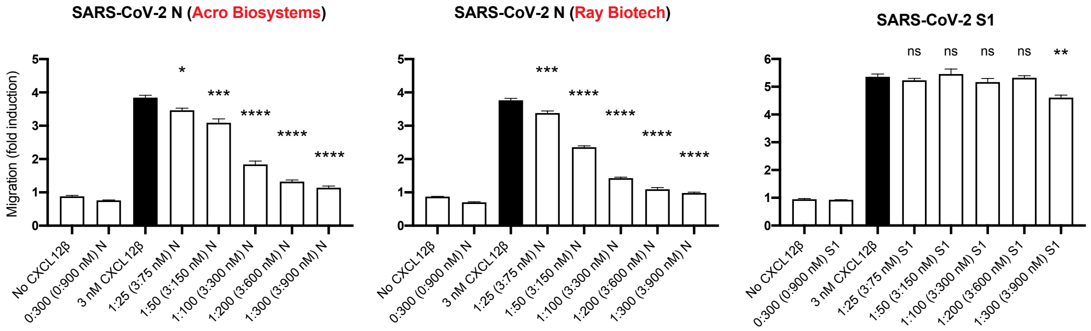


**Supplementary Fig. 2. Additional data supporting in vitro inhibition of chemokine-mediated leukocyte migration mediate by SARS-CoV-2 N but not S.** Chemotaxis of MonoMac-1 cells induced by CXCL12β alone or in the presence of an increasing molar ratio of chemokine : viral protein. In all experiments, CXCL12β was incubated alone or in the presence of SARS-CoV-2 N or S1 (subunit 1 of the S protein) in the lower chamber of transwell migration devices. Migrated cells, added at the beginning of each experiment to the top chamber, were detected in the lower chamber at the end of the experiment. The induction of migration shows means +/- SEM (n = 3) from one representative assay performed in triplicate out of at least three independent assays. One-way ANOVA and Dunnett's Multiple comparison test were used to compare all conditions (except no chemokine and viral protein alone conditions) against migration induced by chemokine alone (dark bars): *ns* (nonsignificant) $p > 0.05$, * $p < 0.05$, ** $p < 0.01$, *** $p < 0.001$, **** $p < 0.0001$.

**a**

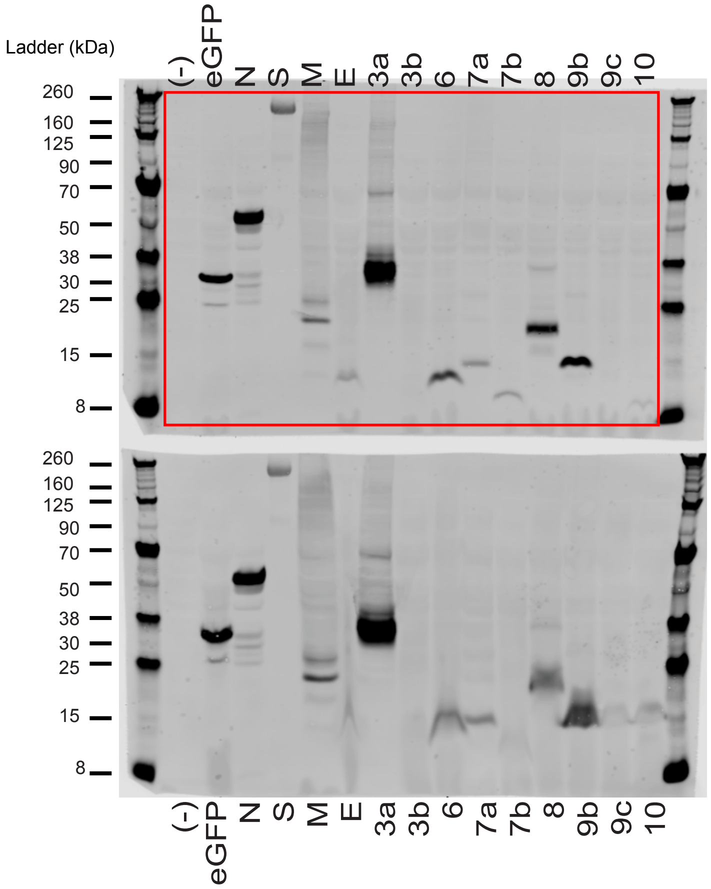


**b**

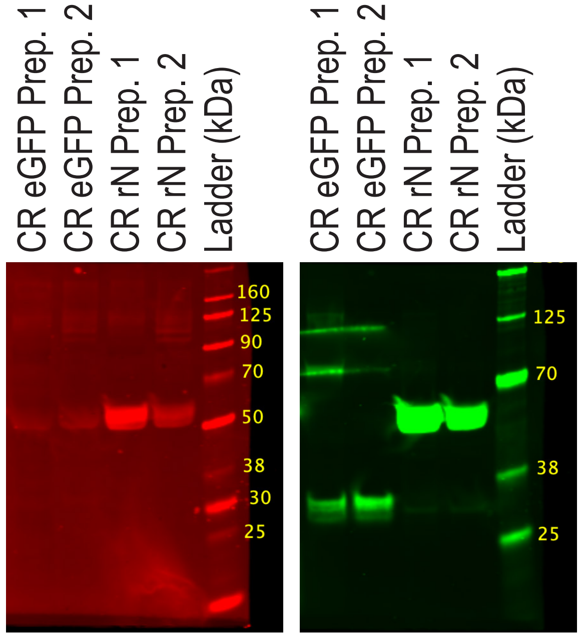


**Supplementary Fig. 3. Immunoblotting data. a,** Unprocessed immunoblot image with molecular weight ladder and red box outlining the image cropping used to generate **Extended Data Fig. 7b**. Each blot corresponds to different protein preparations used for BLI assays. Both blots were stained with primary antibody mouse anti-2xStrep tag (Qiagen #34850, 1:1,000), and secondary goat anti-mouse IgG IRDye 800CW (LI-COR # 926-32210, 1:10,000). **b,** Western blot image with molecular weight ladder from two different preparations of crude lysates (CR, 10 ul loaded) from transfected cells containing recombinant eGFP-2xStrep tag and N-2xStrep tag. The same blot was stained with IRDye 680RD Streptavidin (LI-COR # 926-68079), and human anti-SARS-CoV-2 N mAb (N18) followed by IRDye® 680RD Goat anti-Human IgG Secondary Ab (LI-COR # 926-68078).

### SUPPLEMENTARY INFORMATION LEGENDS

**Supplementary Animations.** This zipped folder contains animations (gif format) generated from maximum intensity projections (MIP) of laser confocal microscopy *z*-stack images of infected cells with wt and SARS-CoV-2_eGFP shown in **Fig. 1, Extended Data Fig. 1 and 2**.

**Supplementary Videos.** This zipped folder contains videos (mov format) generated from sequential laser confocal microscopy *z*-stack images of infected cells with wt and SARS-CoV-2_eGFP shown in **Fig. 1, Extended Data Fig. 1 and 2**. Color code: N (green), S/eGFP (red), DAPI (blue).
