## Supplementary figures and images for "Cell Surface SARS-CoV-2 Nucleocapsid Protein Modulates Innate and Adaptive Immunity"

### 1.Vero cells.gif

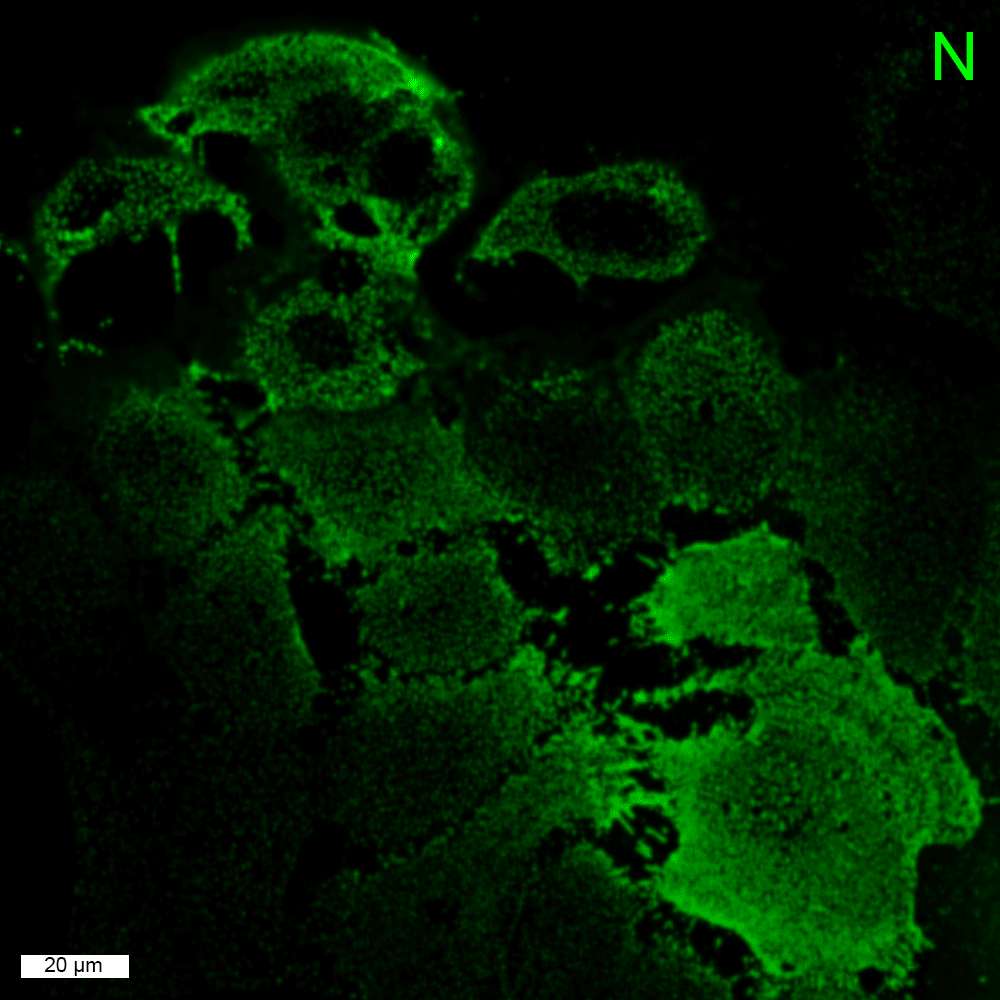

### 1.Vero cells.gif

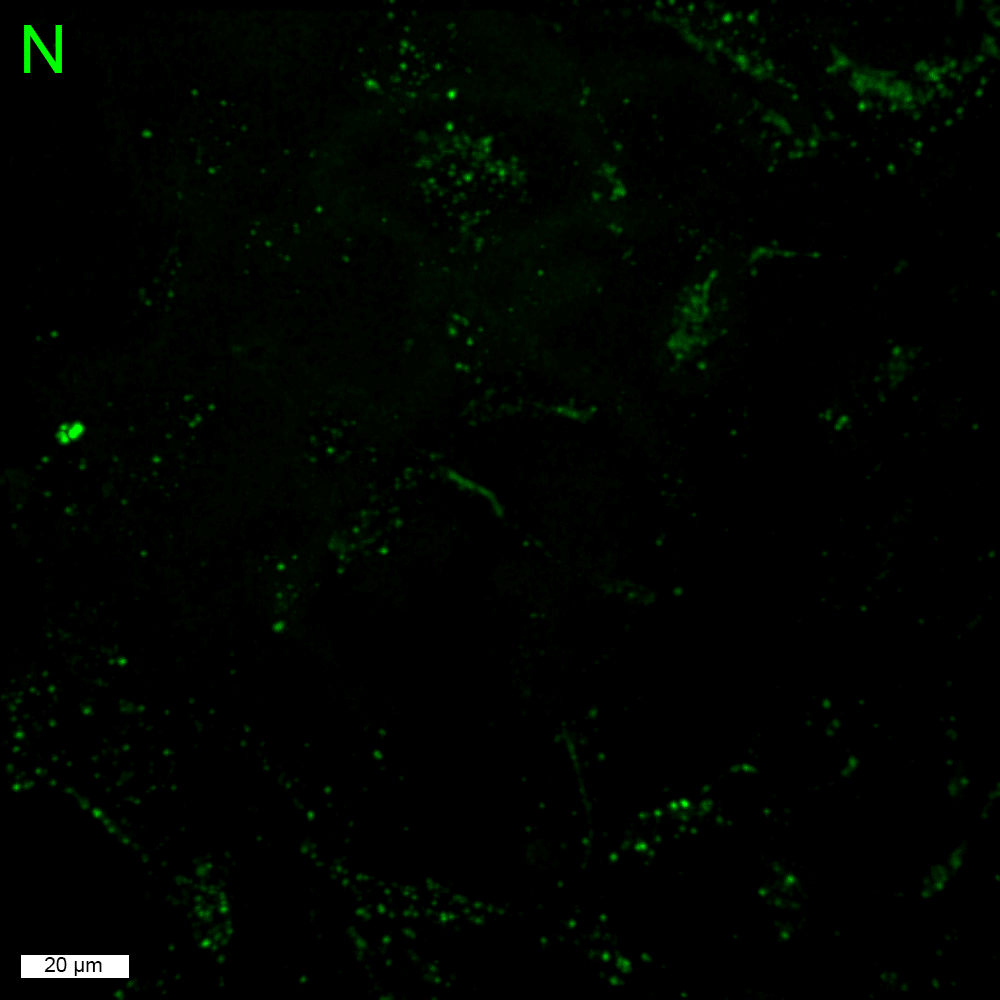

### 2.BHK-21_hACE2 cells.gif

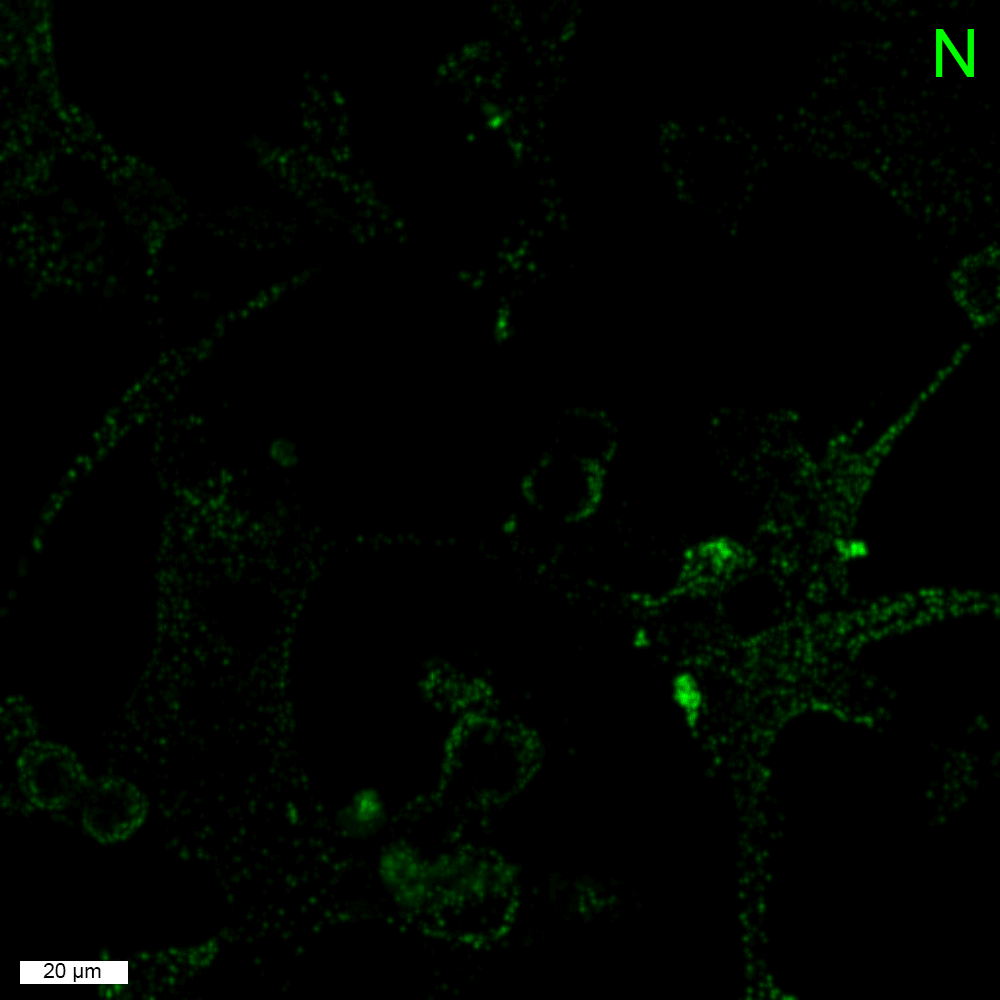

### 2.BHK-21_hACE2 cells.gif

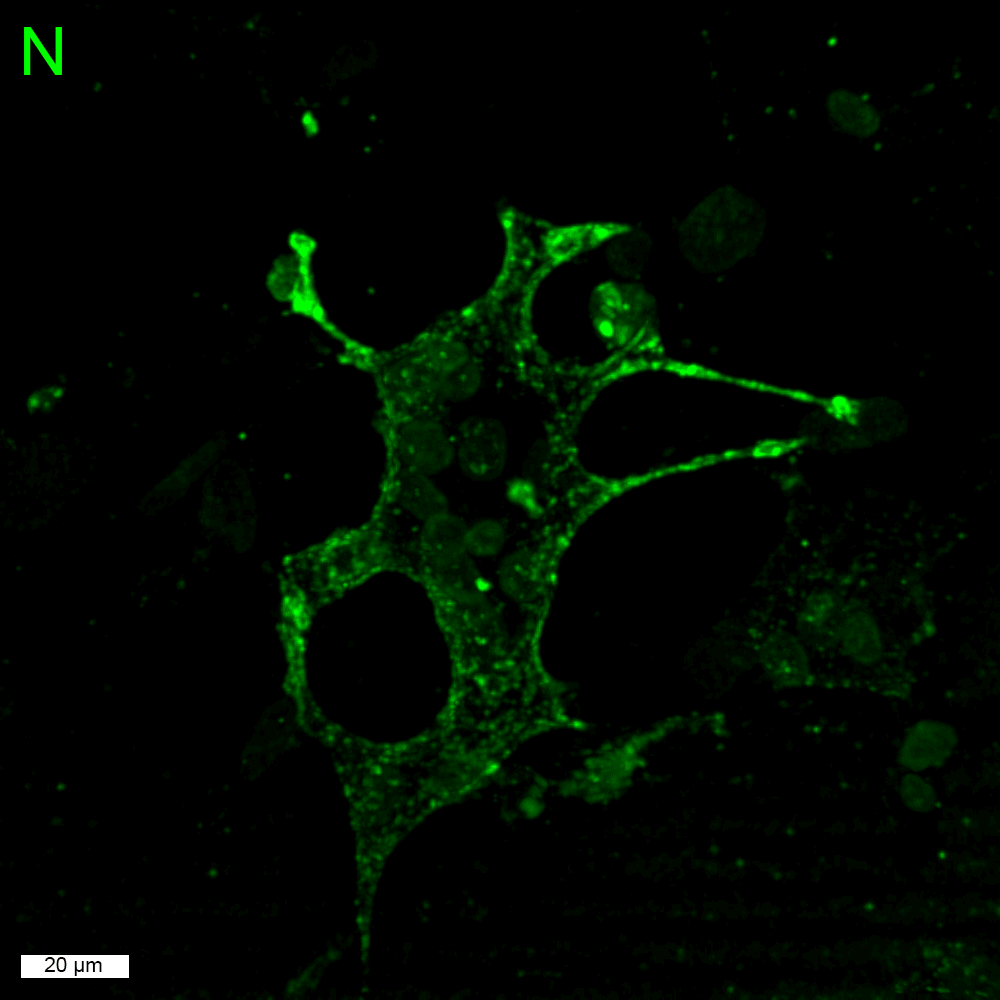

### 3.Caco-2 cells.gif

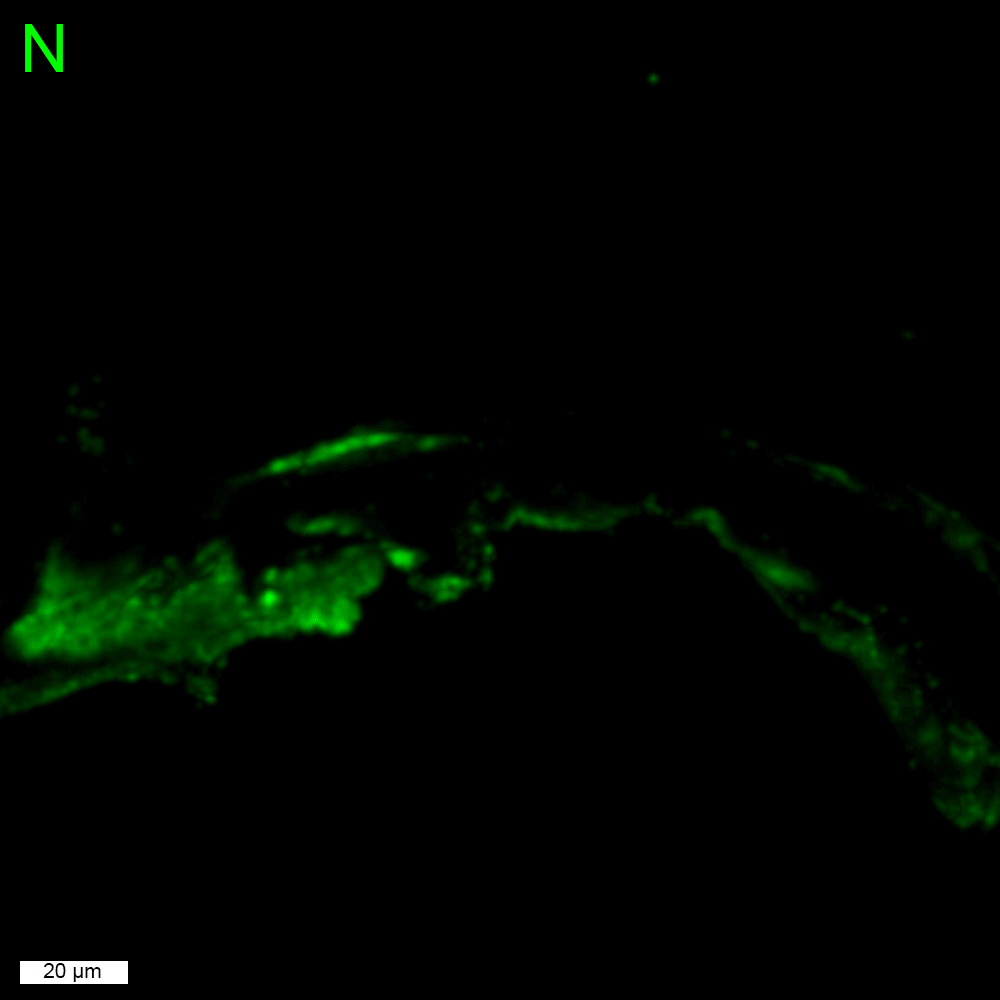

### 3.Caco-2 cells.gif

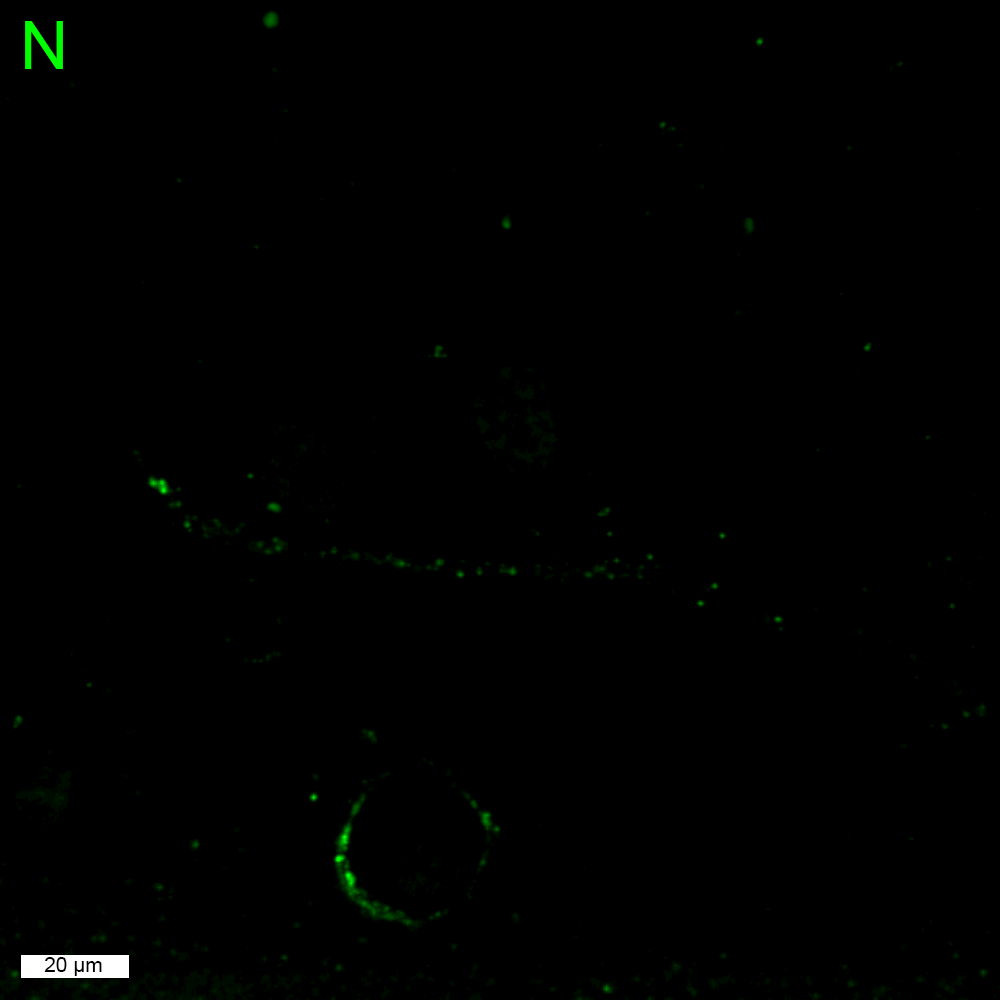

### 4.Calu-3 cells.gif

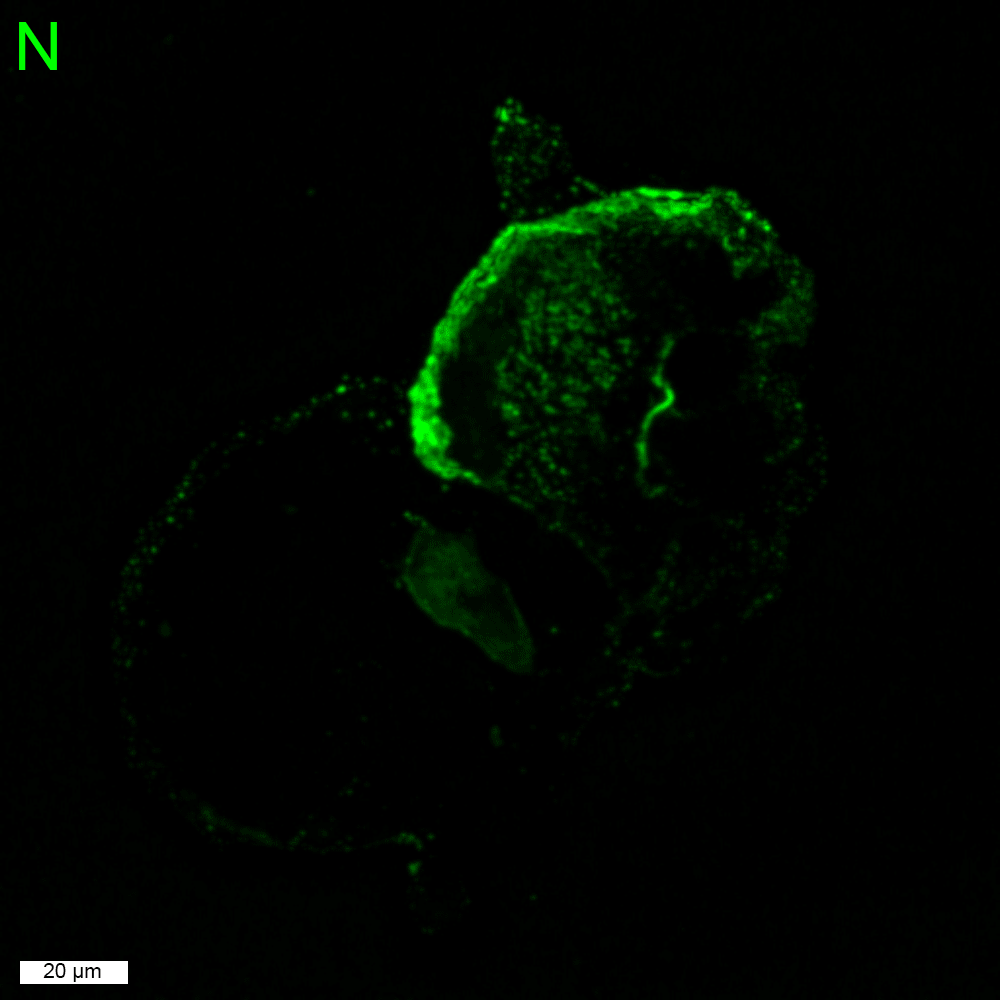

### 4.Calu-3 cells.gif

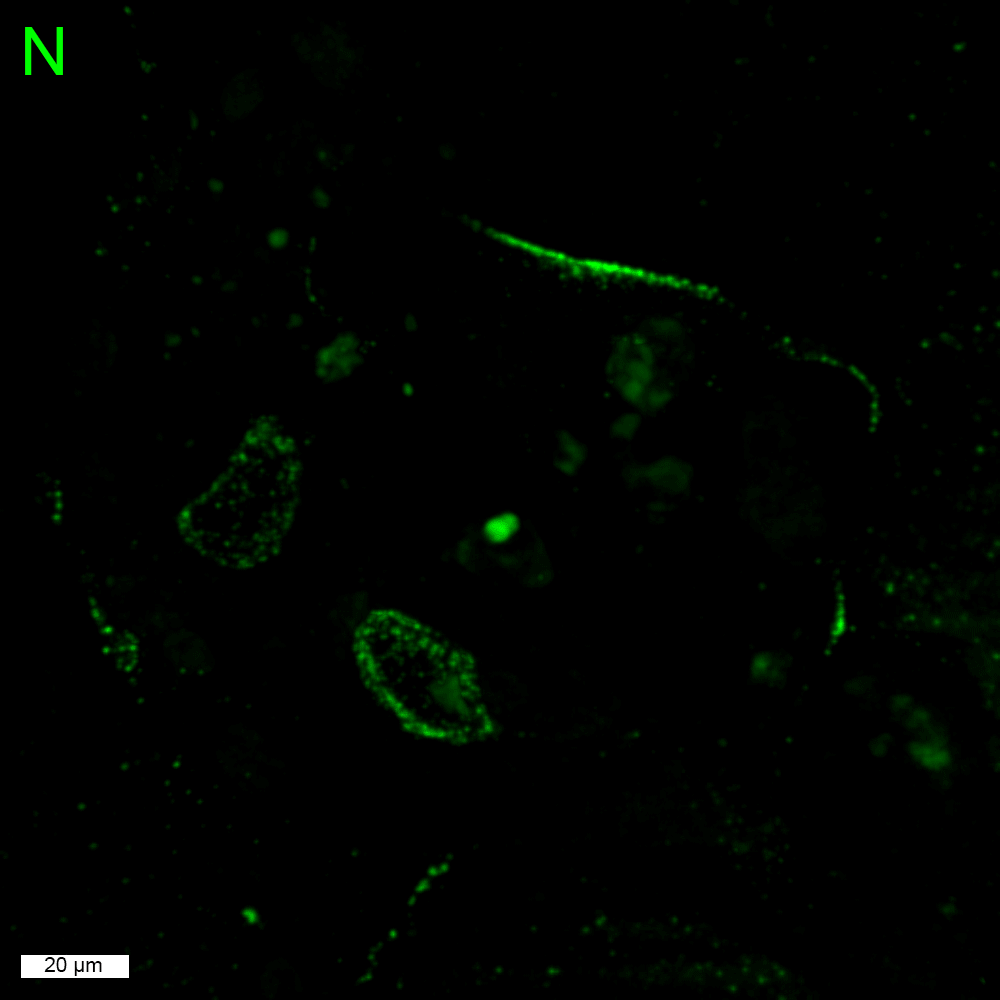

### 5.CHO-K1_hACE2 cells.gif

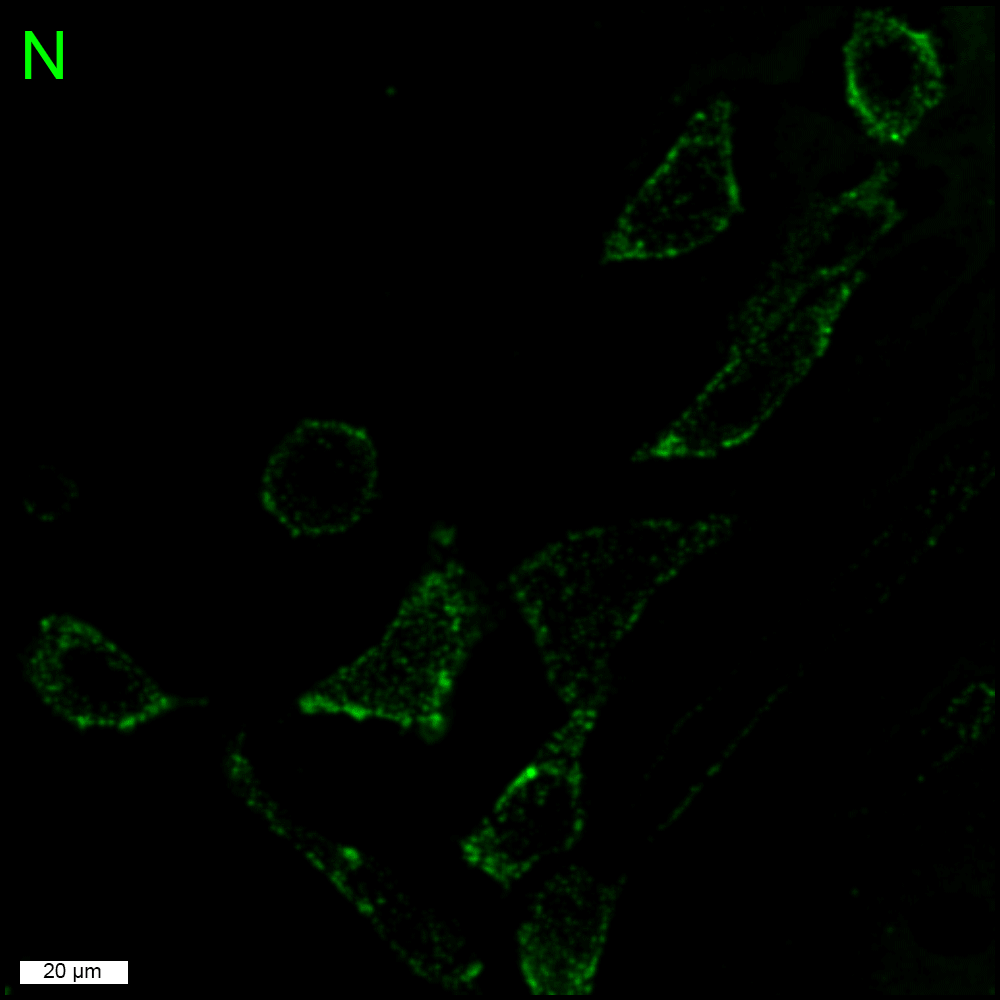

### 5.CHO-K1_hACE2 cells.gif

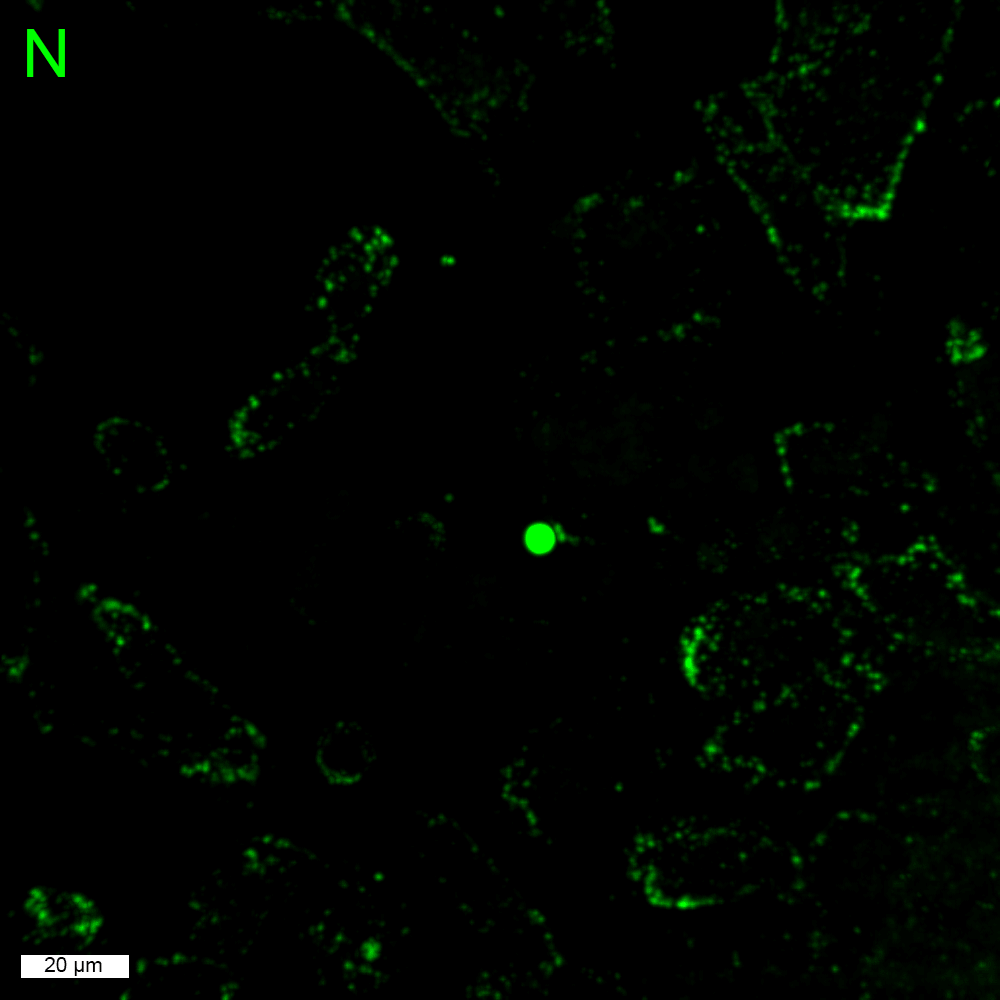

### 6.HEK293-FT_hACE2 cells.gif

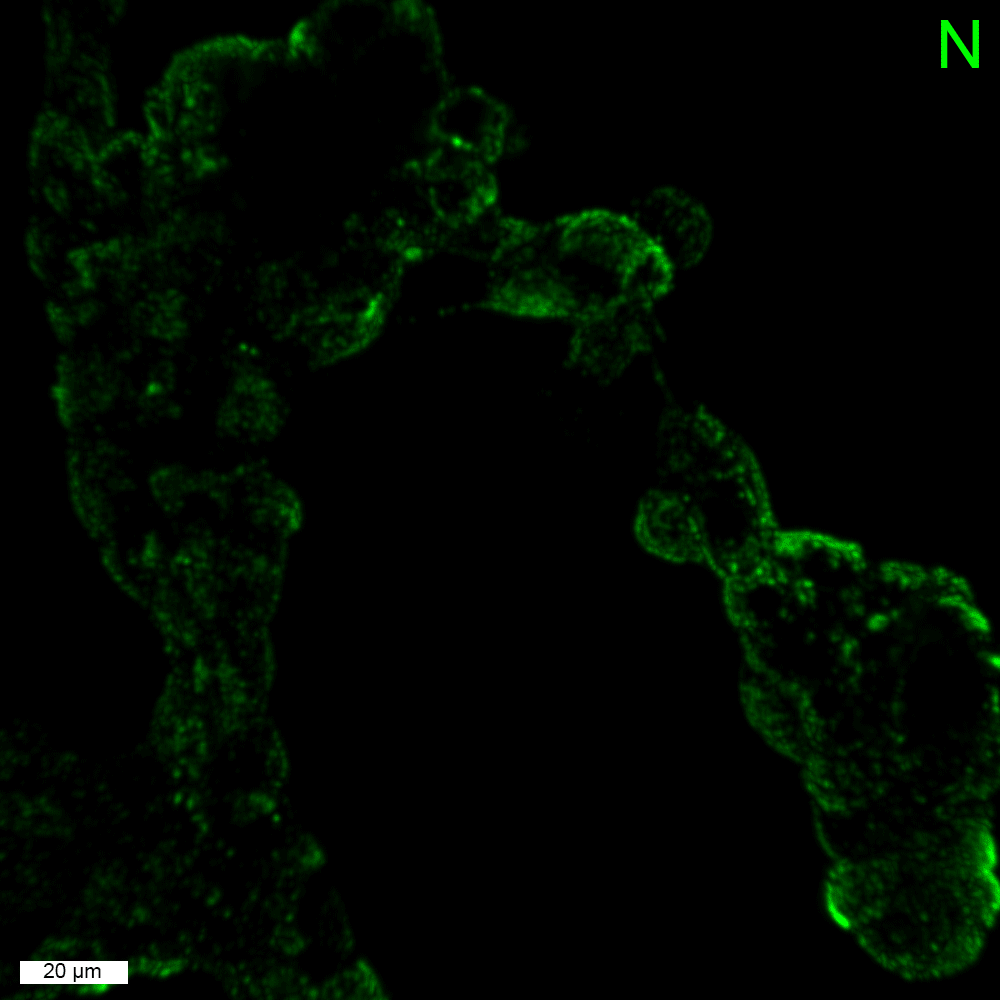

### 6.HEK293-FT_hACE2 cells.gif

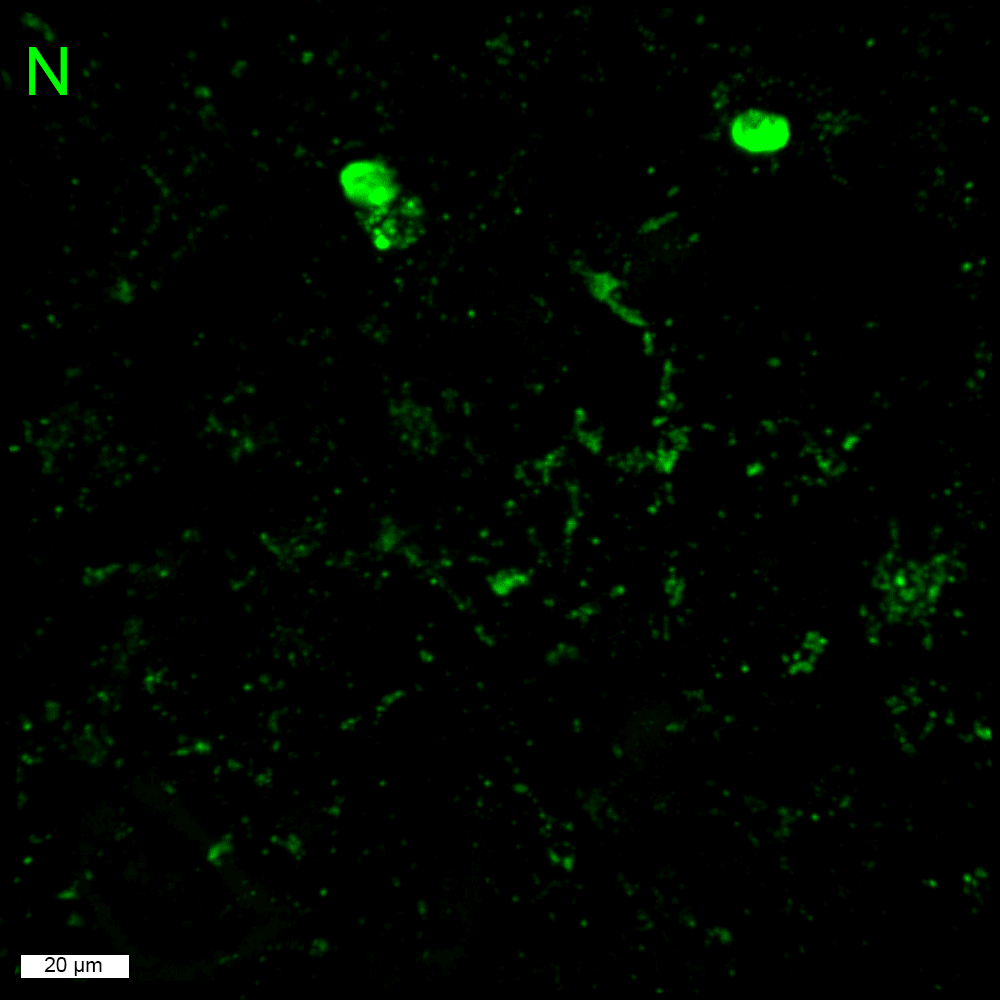

### 7.A549_hACE2 cells.gif

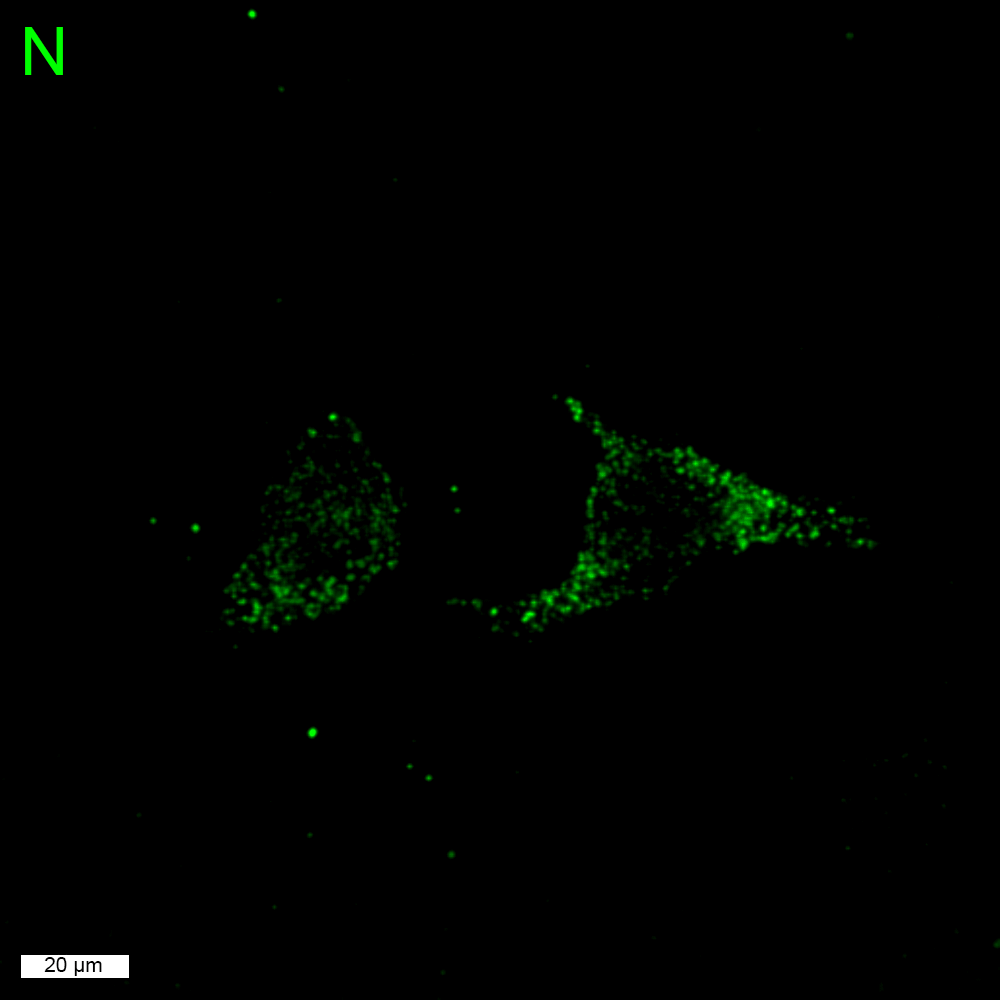

### 7.A549_hACE2 cells.gif

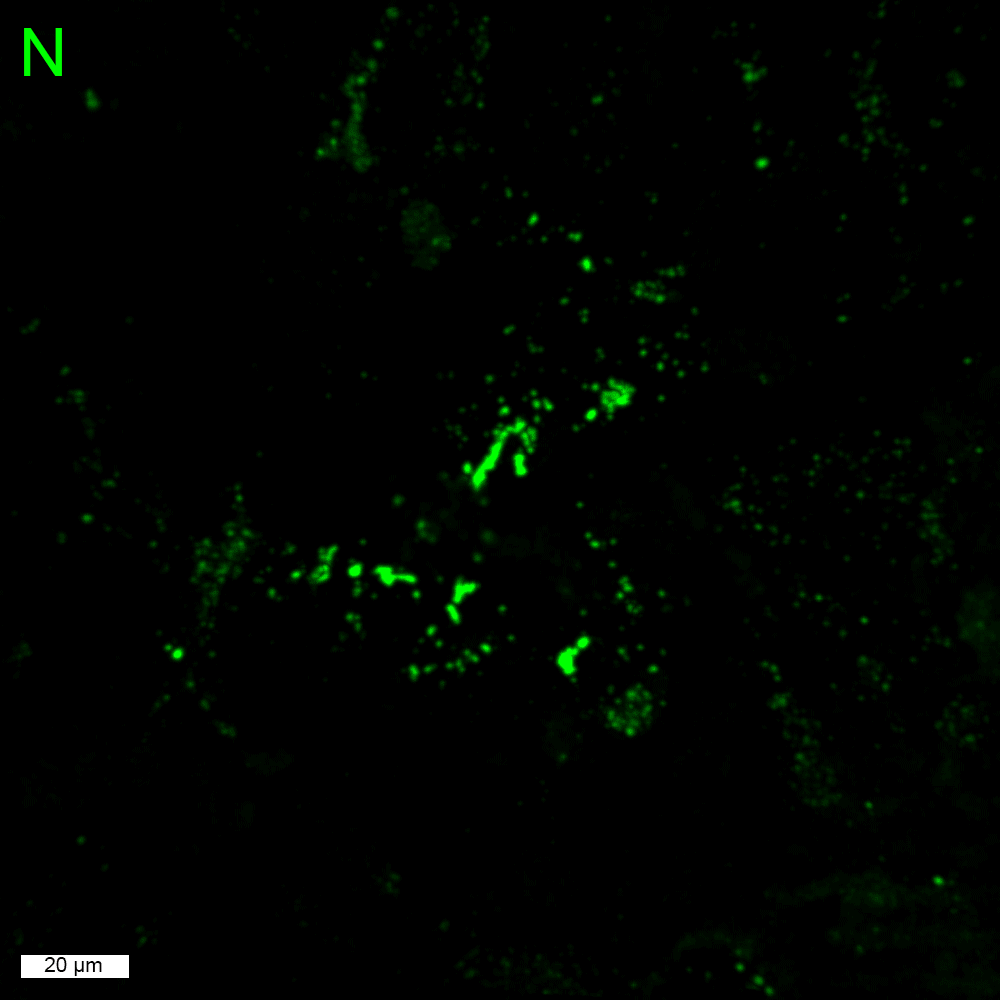
